# Integrative synaptosome multi-omics reveals disrupted synapse organization and localized cryptic transcripts in *C9ORF72*-Frontotemporal Dementia

**DOI:** 10.64898/2026.07.26.740405

**Authors:** Ashton M. Spillman, Eric B. Alsop, Lauren M. Gittings, Krystine Garcia-Mansfield, Ignazio Piras, Anna Bonfitto, Melissa N. Martinez, Ritin Sharma, Kim R. Preller, Matthew Huentelman, Patrick Pirrotte, Kendall Van Keuren-Jensen, Rita Sattler

**Affiliations:** Department of Translational Neuroscience, Barrow Neurological Institute, Phoenix, AZ, USA; Interdisciplinary Graduate Program in Neuroscience, Arizona State University, Tempe, AZ, USA; Bioinnovation and Genome Sciences, Translational Genomics Research Institute, Phoenix, Arizona, USA; Early Detection and Prevention, Translational Genomics Research Institute, Phoenix, AZ, USA; Integrated Mass Spectrometry Shared Resource, City of Hope Comprehensive Cancer Center, Duarte, CA, USA

**Keywords:** Frontotemporal Dementia (FTD), synaptosome, proteomics, transcriptomics, TAR DNA-binding protein 43 (TDP-43), Cryptic Exon (CE)

## Abstract

Frontotemporal Dementia (FTD) and Amyotrophic Lateral Sclerosis (ALS) are linked neurodegenerative diseases characterized by both synaptic dysfunction and TDP-43 pathology. A hexanucleotide repeat expansion (HRE) in the *C9ORF72* (C9) gene represents the most common genetic cause of FTD and ALS, yet the synapse-specific mechanisms underlying disease pathogenesis remain poorly understood. Here, we performed integrated multi-omic profiling of synaptosomes enriched from postmortem frontal cortex and patient-derived induced pluripotent stem cell (iPSC)-derived cortical neurons to define molecular alterations associated with C9-FTD-mediated synaptic dysfunction. Proteomic profiling of frontal cortex-derived synaptosomes identified 1,324 differentially abundant proteins (p<0.05) enriched in pathways regulating synaptic vesicle transport and synapse organization, while synaptosomal RNA sequencing revealed 2,835 differentially expressed protein-coding genes. C9-FTD iPSC-cortical neurons exhibited reductions in excitatory and inhibitory postsynaptic markers, accompanied by progressive impairment of neuronal network activity, supporting both structural and functional deficits. iPSC-derived synaptosomes recapitulated key molecular pathways observed in patient brain, revealing convergent dysregulation of synaptic signaling pathways. Comparative analyses revealed divergence between protein and RNA alterations, consistent with the disruption of regulatory processes that link RNA and protein abundance diseased synapses. Consistent with TDP-43 loss-of-function pathology we identified cryptic exon (CE)-containing transcripts within C9-FTD frontal cortex-derived synaptosomes, including *KALRN* and *STMN2,* providing evidence that aberrantly spliced RNAs localize to synaptic compartments. Together, these findings define convergent molecular pathways underlying synapse vulnerability in both C9-FTD model systems and identify synaptic localization of CE-containing transcripts as a previously unrecognized feature of TDP-43 proteinopathy.

## INTRODUCTION

Frontotemporal Dementia (FTD) is a progressive neurodegenerative disease characterized by behavioral deficits, executive dysfunction, and language impairments due to the selective degeneration of the frontal and temporal lobes [3]. The mean age of onset is between 45 to 64 years of age [21], with a median survival time of 7-13 years after symptom onset, and currently no disease-modifying treatments exist for individuals with FTD. Approximately 20-25% of FTD cases are caused by pathogenic mutations [21], the most common genetic contributor being a hexanucleotide repeat expansion (HRE) in the *C9ORF72* gene (C9) [14, 20, 51]. The identification of the C9-HRE as the most common genetic cause of both FTD and Amyotrophic Lateral Sclerosis (ALS) established these diseases as part of a clinical and pathological spectrum (FTD-ALS) [13, 40].

Mechanistically, the C9-HRE contributes to pathogenesis through both loss and toxic gain-of-function mechanisms, including haploinsufficiency of the C9ORF72 protein, toxic RNA sequestration into foci and formation of dipeptide repeat (DPR) proteins through repeat associated non-AUG (RAN) translation [17]. A convergent pathological hallmark across the FTD-ALS spectrum is the mislocalization and aggregation of the RNA-binding protein TAR DNA-Binding Protein 43 (TDP-43), a DNA and RNA-binding protein that regulates RNA metabolism, transcription, splicing, and transport [25, 39, 43, 49]. In C9-linked disease, TDP-43 is depleted from the nucleus and mislocalized to the cytoplasm, resulting in loss of RNA regulatory functions [47]. A key consequence of TDP-43 loss-of-function is aberrant RNA splicing, including the de-repression of cryptic exons (CE) within gene transcripts, many of which regulate synaptic biology [4, 38, 44]. Initial studies identified CE inclusion in *UNC13A* and *STMN2* [5, 39], while more recent analyses in ALS/FTD patient tissue and TDP-43-depleted iPSC-derived neurons revealed additional synaptic targets, including *KALRN* [18, 23, 45]. These findings support the emerging concept that TDP-43 loss-of-function preferentially disrupts RNA programs critical for synaptic function.

Synaptic dysfunction and subsequent synapse loss are consistent pathological hallmarks across neurodegenerative diseases, including in FTD [9, 11]. Converging evidence from neuroimaging studies from individuals with FTD and pathological assessments of postmortem brain support the presence of progressive decreased cortical connectivity and synaptic degeneration [27, 41, 56]. To investigate synapse-specific molecular changes, isolated synaptic compartments (synaptosomes) have been widely used for proteomic and transcriptomics profiling studies [16, 22]. Synaptosomal analyses in Alzheimer’s Disease (AD) and C9-ALS have identified disease-associated alterations in synapse composition [33, 34]. While prior studies in C9-ALS have reported regional synaptic abnormalities in motor and pre-motor cortex, the molecular landscape of synaptic dysfunction in C9-FTD remains undefined.

Here, we investigate the molecular composition of the “synaptome” in C9-FTD frontal cortex by generating integrated proteomic and transcriptomic profiles of patient-derived synaptosomes. By incorporating mass spectrometry-based proteomics, we identified 1,324 differentially abundant proteins and 2,835 differentially expressed protein-coding transcripts. These changes converge on pathways associated with synaptic vesicle dynamics, neurotransmitter release, and synapse organization. Further, we identified a previously unrecognized synaptic localization of CE-containing transcripts consistent with TDP-43 pathology.

To model observations in frontal cortex tissue, we characterized synaptic dysfunction phenotypes using C9-FTD patient-derived induced pluripotent stem cell (iPSC)-differentiated cortical neurons. These iPSC-cortical neurons were assessed for synaptic density via immunofluorescent staining, in addition to functional assessments of neuronal excitability using longitudinal multielectrode array (MEA) recordings. C9-FTD iPSC-cortical neurons exhibited reduced excitatory and inhibitory postsynaptic marker puncta and inhibitory synapse density, in addition to progressive reductions in neuronal firing activity. To define the molecular alterations associated with these phenotypes, synaptosomes were isolated from the iPSC-cortical neurons and profiled using integrated proteomic and transcriptomic analyses. These analyses demonstrated that C9-FTD iPSC-derived synaptosomes recapitulate key molecular pathways identified in postmortem frontal cortex tissue-derived synaptosomes, supporting the utility of this model for investigating cell-autonomous synaptic abnormalities.

By integrating proteomic and transcriptomic analyses of *ex vivo* and *in vitro* synaptosomes, this study provides an integrated view of the molecular alterations underlying synaptic dysfunction in C9-FTD. These complementary datasets identify pathways associated with synaptic vulnerability and demonstrate for the first time that TDP-43-associated aberrantly spliced mRNAs are present in human synaptic compartments, further implicating RNA misprocessing in synaptic pathology.

## METHODS

### Postmortem tissue samples

Fresh-frozen, postmortem frontal cortex samples from individuals with *C9ORF72*-linked frontotemporal dementia patients (C9-FTD) and neurologically normal control donors were obtained from the Queen Square Brain Bank (QSBB) for Neurological Disorders at University College London. Tissue collection and distribution were performed under appropriate ethical approval. Disease classification was based on neuropathological diagnoses provided by the QSBB, classified as FTLD-TDP A, with clinical presentations of behavioral variant FTD (bvFTD). Summarized demographic information of the tissues used in this study is provided in Table 1, full case demographics are provided in Supplemental Table 1.

**Table 1.** Summary demographics of postmortem tissue sample-derived synaptosomes used in this study.

| **Affliction** | **N** | **Sex (M/F)** | **Age at disease onset (years±SD)** | **Age at death (years±SD)** | **Disease duration (months ±SD)** | **Tissue PMI (hours±SD)** | **Pathological Diagnosis** | **Clinical Diagnosis** | **Frontal Cortex tissue weight (g)** |
| --- | --- | --- | --- | --- | --- | --- | --- | --- | --- |
| **Control** | 10 | 5/5 |  | 83 ± 9.59 |  | 94.17 ± 36.77 | Low level AD | Control (10) | 64.78 ± 2.99 |
| **C9-FTD** | 10 | 6/4 | 58.70 ± 4.62 | 65.70 ± 5.38 | 84 ± 26.91 | 69.47 ± 26.75 | FTLD-TDP-43 Type A | Picks (3), FTD (1), FTLD (1), bvFTD (1), PFNA (2), FTD-MND (1), AD (1) | 68.60 ± 5.25 |

### iPSC culture and cortical neuron differentiation

De-identified C9-FTD fibroblasts (University College London Hospital) and neurologically normal control donor Peripheral Blood Mononuclear Cells (PBMCs, Cedars Sinai Regenerative Medicine Institute) were reprogrammed into iPSCs and quality controlled at Baylor University and the Cedars-Sinai Biomanufacturing Center (CSBC), respectively. All iPSC lines are quality controlled for positive staining of pluripotency markers: SOX2, OCT3/4, and NANOG. Karyotyping of iPSC lines revealed no chromosomal abnormalities.

iPSC lines were cultured in mTeSR™ Plus Medium (StemCell Technologies, #100-0276) under standard physiological conditions: 37°C, 5% CO_2_, and >95% relative humidity. iPSCs were cultured with daily media changes on tissue culture plates coated with Matrigel Matrix (Corning, #B003T18) and monitored for morphology. iPSC colonies displaying smooth edges and compact morphology were expanded prior to neural progenitor cell (NPC) induction.

iPSCs were passaged into single-cell suspensions using ACCUTASE™ (StemCell Technologies, #07922) and were plated in Neural Induction Medium supplemented with 10 µM ROCK inhibitor (Y-27632; StemCell Technologies, #72302). Media was replenished daily for 11 days, resulting in a confluent monolayer of cells. On day *in vitro* (DiV) 12, cells were split using a StemPro EZPassage Disposable Stem Cell Passaging Tool (Gibco, #23181010) to dissociate cells in a grid-like pattern. Cell suspensions were replated onto Neural Maintenance Medium-filled Matrigel-coated plates, supplemented with 10 µM ROCK inhibitor. Media was replenished daily in NMM from DiV13 to 27, with FGF supplementation daily from DiV 13 to 16 (FGF; StemCell Technologies, #78003).

iPSC differentiation into forebrain excitatory cortical neurons followed a previously established protocol using dual SMAD inhibition [52]. DiV 28 NPC were passaged using ACCUTASE and plated onto 6-well plates coated with Matrigel. Cells were seeded at a density of 1 × 10⁶ cells/mL in NMM supplemented with 10 µM ROCK inhibitor, and media replenished 3 times weekly. At DiV 60, cells were seeded for neuronal maturation in 6-well, 24-well plates with glass coverslips (ASI Round Glass Coverslips, 12mm, #SM032), and MEA plate formats, pre-coated with poly-L-ornithine (Sigma-Aldrich, #P4957) and laminin (StemCell Technologies, #200 − 0117). Cortical neurons were aged to DiV 100 for synaptosome isolations, cell fixation for immunofluorescence staining, and MEA analyses. All cortical neurons were quality controlled for the expression of cortical markers: CTIP2, FOXG1, TBR1, SATB2. Demographic information of the iPSC cohort used in this study is provided in Table 2.

**Table 2.**
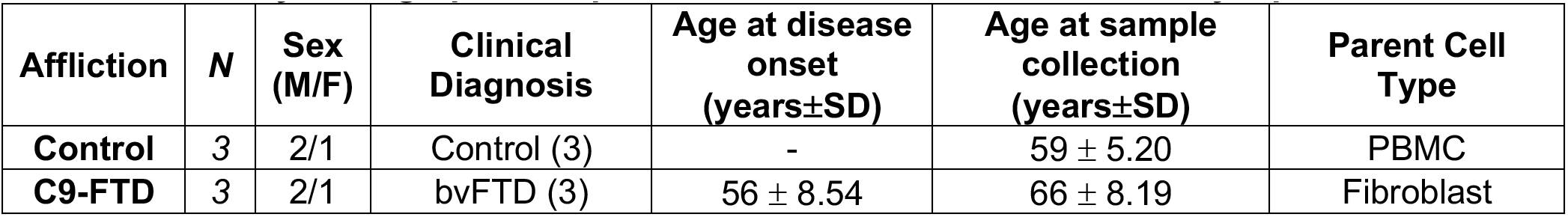
Summary demographics of patient-derived iPSC cortical neuron synaptosomes used in this study.

### Biochemical fractionation and synaptosome isolation

Frozen frontal cortex tissue samples (60-75mg) and 2 million DiV100 iPSC-cortical neurons underwent synaptosome fractionations using a previously published protocol with minor modifications [46]. Briefly, tissue and *in vitro* samples were homogenized with a Teflon douncer, applying 30 strokes in 1mL HEPES-containing Buffer A (10 mM HEPES pH 7.4, 2 mM EDTA, 5 mM Sodium orthovanadate, 30 mM Sodium fluoride, 20 mM β-glycerophosphate and a protease inhibitor cocktail (Roche), pH =7.4). Homogenized lysate (H0) was centrifuged at 500×*g* for 5min at 4°C to pellet cell debris, where the following supernatant was collected and further centrifuged at 10,000×*g* for 15min at 4°C to obtain a crude synaptosome (P2) pellet. The P2 pellet was resuspended in 200 µL Buffer B (50 mM HEPES pH 7.4, 2 mM EDTA, 2 mM EGTA, 5 mM Sodium orthovanadate, 30 mM Sodium fluoride, 20 mM β-glycerophosphate, 1% Triton X-100 and a protease inhibitor cocktail (Roche), pH =7.4). Each P2 fraction was split for protein and for RNA extraction. All fractions were snap-frozen in liquid nitrogen and stored at −80°C.

### Transmission Electron Microscopy (TEM) of Frontal Cortex-derived Synaptosomes

Synaptosome pellets (P2) were isolated from frontal cortex tissue as described above. For transmission electron microscopy, the final P2 pellet was resuspended in 1× PBS containing 2.5% glutaraldehyde to preserve synapstosome ultrastructure for imaging, rather than being solubilized for RNA and protein analyses. Fixed synaptosomes were washed three times with 0.1M Cacodylate buffer, pH=7.2, post-fixed with 1% OsO4 in 0.1M Cacodylate buffer for 30 min and washed three times with 0.1M Cacodylate buffer. The samples were then dehydrated with 60%, 70%, 80%, 95%, 100% ethanol, propylene oxide, and were left in propylene oxide/Eponate (1:1) overnight at room temperature. The next day the vials were left open for 3 hours to evaporate the propylene oxide. The samples were then infiltrated with 100% Eponate and polymerized at ∼64°C for 48 hours. Ultra-thin sections (∼70nm thick) were cut using a Leica Ultra cut UCT ultramicrotome with a diamond knife, picked up on 150 mesh copper EM grids. Grids were stained with 2% uranyl acetate for 10 minutes followed Reynold’s lead citrate staining for 1 minute. Electron microscopy was done on an FEI Tecnai 12 Transmission Electron Microscope equipped with a Gatan OneView CMOS camera.

### Western Blot for Biochemical Synapse Enrichment

Protein lysates derived from biochemical fractionation of brain tissues and iPSC-cortical neurons (H0, P2 fractions) were collected in a triton-containing buffer. Protein concentration of samples was calculated using a bicinchoninic acid (BCA) assay. Western Blot samples were prepared with 5µg of protein mixed with 4x Laemmli Buffer (ThermoScientific, #J60015-AD) and 10x Reducing Agent (Invitrogen, #NP0009), and deionized water. Samples were denatured at 60°C for 10 minutes before loaded onto an SDS-PAGE gel. SDS-PAGE was conducted on 4%-20% Novex Tris-Glycine Plus Midi protein gel, 1mm (Invitrogen, #WXP42026BOX) in 10x Tris-Glycine running buffer. Gels were run at 90 V for 30 minutes and switched to 200 V for 1 hour. Protein transfer was conducted onto 0.2 µm Trans-Blot Turbo Nitrocellulose Membrane (BioRad, #12023955) using the Trans-Blot Turbo system (BioRad), following the manufacturer’s mixed molecular weight transfer protocol. Membranes were stained for total protein using the Revert 700 Total Protein Stain Kit for Western Blot Normalization (LiCOR, #926-11016), imaged, and destained according to the manufacturer’s protocol. Subsequently, membranes were blocked for 1 hour at room temperature in LiCOR Intercept (TBS) Blocking Buffer (LiCOR, #927-60003). Primary antibodies (anti-Synaptophysin, Abcam #16659, 1:200; anti-Homer1, Abcam #184955, 1:500; anti-PSD-95, Abcam #18258, 1:500) were diluted in Intercept T20 (TBS) Antibody Diluent (LiCOR, #927-65003) and incubated overnight at 4°C. Membranes were then washed 3x for 5 minutes each in tris-buffered saline with 1% Triton-X (TBS-T), then incubated for 1 hour at room temperature with HRP-conjugated secondary antibodies (HRP, Donkey anti-Rabbit IgG (H+L) Secondary Antibody; Invitrogen, #31458), and washed 6x for 5 minutes each in TBS-T. SuperSignal West Pico PLUS Chemiluminescent Substrate (ThermoFisher, #34580) was added to the membranes for 5 minutes covered from light at room temperature. Following this reaction, blots were imaged on the Odyssey Imaging System (LiCOR, Odyssey Fc). Band intensities and total protein were quantified and normalized using Empiria Studio Software (LiCORbio v3.3).

### Proteomics Sample Preparation, Data Acquisition and Analysis

SDS was added to each sample to a final concentration of 2%. Samples were lysed using a tip sonicator and clarified by centrifugation at 15,000×*g* for 15 minutes. Protein concentration was quantitated using the Pierce BCA Protein Assay Kit (ThermoScientific, #23225). 25 µg and 5 µg of total protein from brain tissue and iPSC samples, respectively, and were reduced and alkylated with DTT (5 mM final concentration) for 30 minutes at room temperature, followed by iodoacetamide (20 mM final concentration) for 30 minutes in the dark, at room temperature. An SP3 protocol using Sera-mag beads at a protein ratio of 10:1 was employed as previously described for sample clean-up [28]. After clean-up, samples were digested overnight on-beads using trypsin endoproteinase (Trypsin Gold, Promega) at an enzyme:substrate ratio of 1:25. Following digestion, samples were acidified with 100% formic acid, placed on a magnetic rack and the supernatant containing digested peptides was collected. The beads were washed twice more with 50 µL of 50 mM ammonium bicarbonate and the resulting supernatant was pooled with the eluted peptides. Peptides were vacuum dried and reconstituted in LC-MS grade water containing 0.015% dodecyl-maltoside and 0.1% fomic acid.

Proteomics data were acquired on a NanoElute 2 liquid chromatography system coupled to a timsTOF Ultra 2 mass spectrometer (Bruker Daltonic). For each sample, 50 ng tryptic peptides in 3 µL injection volume were directly loaded on a C18 column (Ionopticks Aurora Ultimate, 25 cm length, 75 µm ID, 1.7 µm particle size and PepSep Ultra, 25 cm length, 75 µm ID, 1.5 µm particle size for tissues and iPSCs, respectively). Peptides were eluted using a 30-minute reverse phase chromatography gradient formed by Solvent A (LC-MS grade water, 0.1% formic acid) and Solvent B (LC-MS grade acetonitrile, 0.1% formic acid) at a flow-rate of 250 nL/min. Data acquisition was performed in diaPASEF mode using a mass range of 300-1000 m/z covered by 28 MS/MS windows with a cycle time of 0.96 second. Capillary voltage was kept at 1600 V and dry gas temperature was set to 200°C.

For brain tissue samples, raw spectra were searched against a human protein fasta (UP000005640, downloaded December 2025) in directDIA+ mode through Spectronaut (v20.2). Default settings were used with the exception that cross-run normalization and imputation were not selected. In short, theoretical trypsin digestion allowed up to 2 missed cleavages. Carbamidomethylation (C) was set as a fixed modification, with acetylation (N-term) and oxidation (M) as variable modifications. PSM, peptide, and proteins were filtered for FDR <0.01. Protein scoring used the Highest Scoring Observation method to align with version 19 of Spectronaut. For iPSC samples, proteins and peptides were identified using Spectronaut (v19.7) in directDIA+ mode. Raw spectra were mapped against a SwissProt/UniprotKB human fasta file (UP000005640, downloaded January 2025), with trypsin digestion allowing for up to 2 missed cleavages. Cysteine carbamidomethylation was set as a fixed modification, with methionine oxidation and N-term acetylation as variable modifications. PSMs, peptides, and proteins were filtered for FDR <1%. Protein abundances were normalized via variance stabilizing normalization (VSN) in R (v4.1.2). Statistical significance was calculated using a Welch’s t-test, followed by a Benjamini-Hochberg correction for multiple comparisons, and determined using nominal p<0.05.

### RNA Isolation, Library preparation, and Sequencing

RNA from homogenate and synaptosome fractions from both frontal cortex and iPSC-cortical neuron samples were extracted using the RNeasy Plus Micro Kit (Qiagen, #74034), following the manufacturers protocol. RNA was quantified fluorometrically using the Qubit Assay (Invitrogen, #Q33265). A total of 10ng of RNA per sample was treated with the Heat&Run gDNA Removal Kit (ArcticZymes Technologies, #80200-250) and subsequently used for library preparation with the SMARTer Stranded Total RNASeq Pico Input v3 Kit (Takara Bio Inc, #634487), following the manufacturers protocol, with a final elution volume of 32µL. Library quality and fragment size distribution were assessed using the TapeStation High Sensitivity D1000 Assay (Aglient Technologies, #5067-5584), selected due to higher final elution volume. Libraries were pooled to a final concentration of 1.1nM and spiked with 1% PhiX control (Illumina). Sequencing was performed on an Illumina NovaSeq X platform using a 10B flow cell with paired-end reads (151–9–9–151 cycle configuration) after rebalancing libraries with iSeq data.

### Transcriptomic Data Analysis

Fastq files were generated from raw sequence files using bcl2fastq (Illumina) using default parameters. Reads were trimmed with cutadapt v1.17 according to kit recommendations. Trimmed fastq files were then aligned to the GRCh38 genome with STAR v2.6.1d with the following parameters: --runMode alignReads --outSAMtype BAM Unsorted --outSAMmode Full --outSAMstrandField intronMotif --outFilterType BySJout --outSAMunmapped Within --outSAMmapqUnique 255 --outFilterMultimapNmax 20 --outFilterMismatchNmax 999 --outFilterMismatchNoverLmax 0.1 --alignMatesGapMax 1000000 --seedSearchStartLmax 50 --alignIntronMin 20 --alignIntronMax 1000000 --alignSJoverhangMin 18 –alignSJDBoverhangMin 18 --chimSegmentMin 18 --chimJunctionOverhangMin 18 --outSJfilterOverhangMin 18 18 18 18 --alignTranscriptsPerReadNmax 50000. Following genome alignment, reads were counted with featureCounts v1.6.3, (part of the subread package) using a non-redundant genome annotation combined from GENCODE 47 and LncBook v2.1 and the following parameters: -p -t exon -g gene_id. Additionally, the strandedness parameter was passed to featureCounts. Raw gene count tables for use with downstream analyses were generated using a python script that combined all featureCounts output files into a single TSV file.

### Cryptic Exon Junction Analysis

Junctions were identified as described in [18], following [5]. Briefly, fastq files were aligned to the human genome (GRCh38) using STAR and standard alignment parameters. Resulting BAM files were then analyzed with regtools junctions extract (options: -a 6 -m 30 -M 500000) to generate junction files, which include novel junctions, for all samples [10]. Leafcutter’s leafcutter_cluster_regtools.py (options: -m 10 -p 0.0001) was then used to generate a summary counts matrix of all junctions found in the dataset [36]. Counts of junctions of interest across samples were then examined from the leafcutter summary output file.

### Gene Ontology Analysis

Gene set over-representation analyses were performed using SynGO (v1.3, 2025) over-representation analysis was performed [32] . Gene sets used for Gene Ontology (GO) Biological Process (BP) terms were selected across datasets for genes that passed significance thresholds (p<0.05) and stratified into upregulated and downregulated terms based on log_2_ fold-change.GO terms displayed were selected from the top 10 terms based on hierarchical p-adjusted value (FDR-corrected p-value) that contained a GO:BP ID number, and the number of genes in the gene set that corresponded to the respective GO:BP term.

### Data Visualization

Data visualization and statistical analyses were performed in RStudio (v2024.12.1+563). Differential gene expression analysis of RNA-seq data was conducted using DESeq2. To ensure consistency in brain and iPSC samples across protein and RNA datasets, a uniform significance threshold of p<0.05 was applied for differential expression and downstream gene set enrichment analyses. Data processing and visualization were performed using packages ggplot2, dplyr, stringr, and paletteer. PCA plots, Volcano plots, and dot plots for gene set enrichment analysis were generated using ggplot2-based workflows. Heatmaps were generated using ComplexHeatmap. Set intersection analyses were visualized using ggVennDiagram and UpSetR. Plot annotation and labeling were facilitated using ggrepel. Genes included in each multi-omic heatmap were selected based on the magnitude of differential expression across the frontal cortex proteomic, frontal cortex transcriptomic, iPSC proteomic, and iPSC transcriptomic datasets. Genes belonging to the intersecting SynGO Biological Process gene sets were ranked using a composite effect size, calculated as the sum of the absolute log_2_ fold-change values across the frontal cortex proteomic, frontal cortex transcriptomic, iPSC proteomic, and iPSC transcriptomic datasets. The top 25 ranked genes were selected for visualization. Additional data visualization and analysis was conducted in GraphPad Prism (v10.4.1), using various comparison tests (Mann-Whitney test, Welch’s t-test, 2-way ANOVA).

### Immunofluorescent staining of Synaptic Markers and Puncta Quantification

On DiV 100, cell culture media was aspirated, and cells were washed in phosphate-buffered saline (PBS) before methanol fixation in 100% methanol for 5 minutes at 4°C. Following fixation, cells were washed in PBS and permeabilized and blocked in a 5% Donkey Serum-containing buffer with 0.25% Triton-X in PBS for 1 hour at room temperature. Cells were incubated with primary antibodies for synaptic and neuronal markers (Supplemental Table 2) diluted in blocking solution overnight at 4°C. After 3 PBS washes, cells were incubated with fluorescently conjugated secondary antibodies (Supplemental Table 2) for 1 hour in the dark at room temperature. Following 3 PBS washes, cells were stained with DAPI (Invitrogen, #D1306) for 7 minutes to label nuclei. Cells were washed 2 additional times in PBS before mounting coverslips to slides using ProLong Glass Antifade Mountant (Invitrogen, #P36980). Synaptic markers were quantified from 10 randomly selected fields of view on a 63x objective (Zeiss), providing robust sampling across technical replicates per differentiation. Each image was captured using a z-stack of 5-6 images at intervals of 0.5 µm. Confocal data was analyzed for synaptic puncta by first identifying neuronal surface filaments of MAP2-positive cells, and the circularity of synaptic puncta (0.3µm to 0.5µm) in Imaris (v10.1.1), specialized for the detection of particles. Puncta counts were normalized against the area of MAP2 for the density of dendritic networks, based on [55]. Synaptic puncta were determined based on sphericity and determined to be localized onto dendrites based on proximal distance to the MAP2 signal within 1µm. The background signal outside of MAP2 selection was subtracted. Colocalization quantification was determined by proximity of presynaptic and postsynaptic puncta, within one radius of the proximal puncta. Values were compared using Mann-Whitney Test in GraphPad Prism (v10.4.1) to test for differences between control and disease groups.

### Microelectrode Array (MEA) recordings of iPSC-cortical neurons

Neurons were seeded at 50,000 cells per well of a 48-well CytoView MEA 48 (Axion Biosystems, M768-tMEA-48B) plate at DiV 60. MEA recordings commenced 5 days post seeding to allow for cells to settle. MEA recordings were conducted on the Maestro Middleman (MAESTRO768D, #1000589) once weekly prior to media changes, at the same time of day. Neurons were equilibrated in the recording chamber for 5 minutes prior to the start of recording. Neuronal cultures were recorded for 5 minutes. Metrics shown in this study are from 5-minute recordings. Standard filters were applied across recordings, including Butterworth high pass (200Hz) and low pass (3000Hz) filters, and 5.5 standard deviations (SD) of signal-to-noise ratios. Data presented shows the average metric (±SEM) of each condition across each cell line derived from control donor or C9-FTD patient. Values were compared with a Welch’s T-test or 2way ANOVA in GraphPad Prism (v10.4.1) to test for differences between control and disease groups.

## RESULTS

### Successful enrichment of synaptic proteins and RNAs in isolated brain synaptosomes

Synaptic dysfunction is a pathological hallmark of neurodegenerative diseases, including C9-FTD. To define synapse-specific molecular alterations in C9-FTD, we isolated synaptosomes from postmortem frontal cortex tissue obtained from clinically and pathologically confirmed C9-FTD cases and age-matched neurologically normal controls (n = 10/group; Table 1), as well as from iPSC-derived cortical neuron cultures generated from individuals with C9-FTD and healthy controls (n = 3/group). Isolated synaptosome fractions were subjected to mass spectrometry-based proteomic and RNA sequencing analyses (Fig. 1a). Successful enrichment of synaptic compartments was confirmed by western blot detection of presynaptic (Synaptophysin, SYP) and postsynaptic (Homer1) marker proteins within P2 fractions (Supplement Fig.1a,b). To assess the structural integrity of the isolated synaptosomes, TEM was performed on frontal cortex-derived synaptosome preparations from control and C9-FTD cases. Representative P2 preparations contained membrane-enclosed presynaptic terminals with synaptic vesicles and electron-dense postsynaptic material, consistent with intact synaptosomes (Fig. 1b).

**Figure 1.**
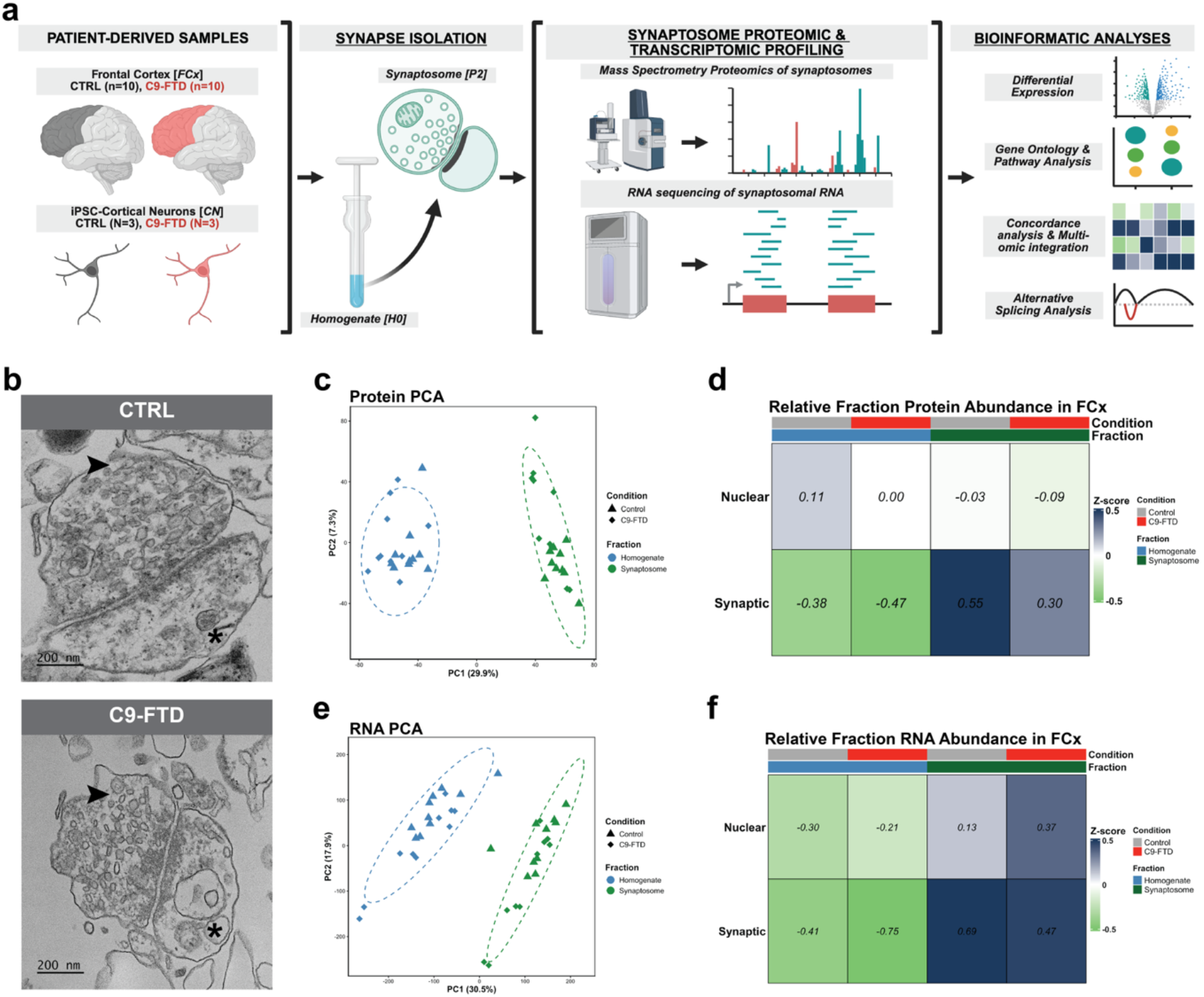
Multi-omic Profiling of Frontal Cortex-Derived Synaptosomes in C9-FTD. **a: Experimental Workflow:** Schematic diagram illustrating the workflow of synaptosome isolation, mass spectrometry and RNA sequencing and analyses from frontal cortex tissue samples from subjects with C9-FTD (n=10) and age-matched controls (n=10), in addition patient-derived iPSC-cortical neuron models from C9-FTD (N=3) and non-neurological disease controls (N=3). **b: Ultrastructural Validation:** Representative Transmission Electron Micrographs of isolated synaptosomes from control (top) and C9-FTD (bottom) frontal cortex tissue. *Arrow indicates presynapse; asterisk indicates postsynapse*. *Scale bar=200nm; magnification=42,000x.* **c: Proteomic separation of homogenate and synaptosome fractions:** Principal Component Analysis (PCA) of proteomic data from homogenate (H0) and synaptosome (P2) fractions (PCA1=29.9%, PC2=7.3%), demonstrating sample separation by biochemical fraction identity. **d: Enrichment of Synaptic and Nuclear Markers:** Z-score-normalized heatmap of synaptic protein abundance across H0 and P2 fractions in control and C9-FTD cases, highlighting enrichment of synaptic marker genes in the P2 fraction and nuclear marker gene enrichment in the H0 fraction. **e: Transcriptomic separation of homogenate and synaptosome fractions:** PCA of transcriptomic data from frontal cortex tissue-derived H0 and P2 fractions (PC1=30.5%, PC2=17.9%), demonstrating sample separation by biochemical fraction identity. **f: Enrichment of Synaptic and Nuclear Markers:** Z-score-normalized heatmap of synaptic RNA abundance across H0 and P2 fractions, highlighting enrichment of synaptic marker genes in the P2 fraction. **d, f) Fraction Purity Validation:** Heatmaps showing Z-score normalized abundance of synaptic and nuclear markers. The Synaptosome fraction shows a strong positive enrichment of synaptic markers compared to the Homogenate. Conversely, nuclear markers are depleted in the Synaptosome fraction relative to the Homogenate, confirming successful separation of cytoplasmic synaptic terminals from nuclear contaminants.

Principal Component Analysis (PCA) of the proteomic data from frontal cortex revealed clear separation between homogenate (H0) and synaptosome (P2) fractions, with fraction identity representing the primary source of variance (PC1=29.9%), followed by disease status (PC2=7.3%; Fig.1c). To further validate enrichment of synaptic proteins in the isolated synaptosomes, we compared compartment-specific protein abundance across the H0 and P2 fractions. Consistent with successful subcellular enrichment, synaptic proteins defined by the SYNGO database [32] were enriched in P2 fractions, whereas H0 fractions preferentially contained nuclear-associated marker proteins, defined by the MGI GO database [15] (Fig.1d). Consistent with the protein-level observations, similar separation between H0 and P2 fractions was observed in the transcriptomic datasets (PC1=30.5%, PC2=17.9%; Fig.1e), with synaptic transcripts enriched within isolated synaptosomes (Fig.1f).

Together, these orthogonal structural, biochemical, proteomic, and transcriptomic analyses demonstrate successful enrichment of synaptic material and establish a robust platform for defining synapse-specific molecular alterations in C9-FTD.

### Synaptosome proteomics reveals synapse-selective molecular changes in C9-FTD

To assess synapse-specific proteomic alteration in C9-FTD, we performed mass spectrometry-based proteomics on frontal cortex homogenates and enriched synaptosomes from control and C9-FTD cases. Comparative analysis across homogenate and synaptosome fractions identified both global and compartment-specific disease-associated changes. Overall, 21.9% of proteins detected in homogenates and 14.5% of proteins detected in synaptosomes were significantly altered in C9-FTD (p<0.05; Fig.2a).

Despite proteomic overlap between homogenate and synaptosome fractions, each fractions exhibited distinct proteomic profiles (Fig.2b). To further assess compartment-specific proteomic alterations, we identified differentially abundant proteins (DAPs; p<0.05) in homogenate and synaptosome fractions, stratified by directionality. An UpSet plot illustrating the overlap of DAPs across fractions revealed a substantial subset of DAPs unique to the synaptosome fraction (Fig. 2c). In contrast, a smaller subset of proteins was consistently altered across both fractions, suggesting broader, disease-associated changes. Notably, several proteins exhibited opposing directions of change between homogenate and synaptosome fractions, consistent with synapse-specific protein redistribution or localized synaptic remodeling in C9-FTD.

**Figure 2.**
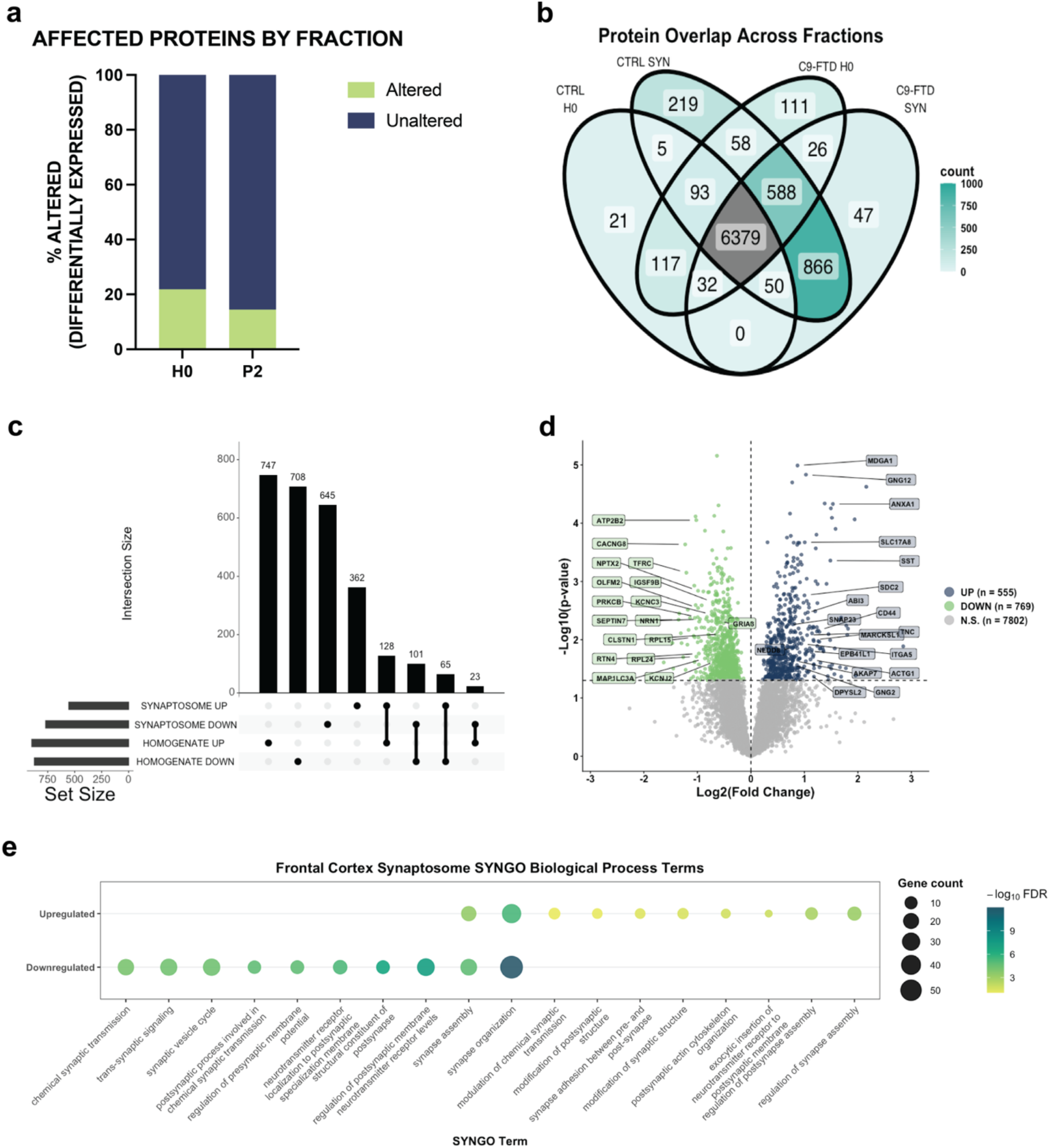
Differential Proteomic Landscape of C9-FTD Frontal Cortex and Synaptosomes. **a: Differentially Abundant Proteins by Fraction:** Differential expression analysis identified disease-associated proteomic changes in both the homogenate and synaptosome fractions, with a greater proportion of quantified proteins altered in the homogenate (21.9%) than in the synaptosome fraction (14.5%). **b: Overlap of proteins detected across H0 and P2 fractions**: Venn diagram displaying the distribution of proteins identified across H0 and P2 fractions in control and C9-FTD cases. **c: Intersection of Differentially Abundant Proteins (DAPs):** UpSet plot displaying the overlap of DAPs across fractions. The high number of unique synaptic DAPs (vertical bars with single dots) suggests that C9-FTD pathology is highly compartmentalized. Conversely, the intersection of Homogenate and Synaptosome DAPs identifies a subset of proteins with consistent global dysregulation, as well as those showing opposite directions of change across compartments. **d: Synaptosome Differential Expression:** Volcano plot of DAP identified in C9-FTD synaptosomes (n=1,324), highlighting the top 20 synaptic proteins annotated based on absolute log_2_FC and SYNGO annotation. A total of 555 proteins were significantly upregulated, and 769 proteins were downregulated (p=<0.05, *t-test*). **e: SYNGO Biological Process Enrichment:** SYNGO Biological Process enrichment analysis of significantly upregulated and downregulated proteins in synaptosome fractions, displaying the top 10 enriched unique and shared terms per direction (8 unique and 2 shared) (p=<0.05, relative to C9-FTD).

To further investigate synapse-specific proteomic alterations in C9-FTD, we next focused on the synaptosome proteome. Proteomic profiling of synaptosomes identified 1,324 DAPs in C9-FTD, including 555 upregulated and 769 downregulated proteins relative to controls (p<0.0; Fig.2d). Functional annotation of the most highly altered synaptic proteins, ranked by log_2_ fold change, using the SYNGO database highlighted widespread dysregulation of proteins associated with excitatory and inhibitory synapses, including pathways involved in vesicle trafficking and postsynaptic organization. These findings suggest extensive proteomic remodeling at both presynaptic and postsynaptic compartments in C9-FTD.

To determine which synaptic functions were most affected at C9-FTD synapses, SYNGO Biological Process (BP) enrichment analysis was performed on upregulated and downregulated proteins. Upregulated proteins demonstrated modest enrichment for pathways related to synaptic structural remodeling and synapse organization, including SYNGO BP terms: *regulation of synapse assembly; regulation of postsynapse assembly; synapse adhesion between pre- and post-synapse; modification of synaptic structure; modification of postsynaptic structure; postsynaptic actin cytoskeleton organization; exocytic insertion of neurotransmitter receptors into the postsynaptic membrane; modulation of chemical synaptic transmission*. In contrast, downregulated proteins were strongly enriched for pathways related to synaptic signaling and neurotransmission, including SYNGO BP terms: *synaptic vesicle cycle; chemical synaptic transmission; trans-synaptic signaling; postsynaptic processes involved in chemical synaptic transmission; regulation of presynaptic membrane potential; regulation of postsynaptic membrane neurotransmitter receptor levels; neurotransmitter receptor localization to postsynaptic specialization membrane; structural constituent of postsynapse*. Terms associated with *synapse organization* and *synapse assembly* were enriched in both upregulated and downregulated datasets, suggesting widespread remodeling of the synaptic architecture in C9-FTD.

### Synaptosomal transcriptomics reveal widespread RNA dysregulation and distinct transcriptomic–proteomic alterations in C9-FTD

To characterize the synaptic transcriptome in C9-FTD, we performed RNA sequencing of frontal cortex-derived synaptosomes from C9-FTD and neurologically normal control cases. Analysis of RNA biotypes demonstrated that protein-coding transcripts represent the majority of RNA species (>60%) in both control and disease synaptosomes, with additional contributions from long non-coding RNA (lncRNA; Fig.3a). Overall, RNA biotype distribution was largely preserved between groups.

**Figure 3.**
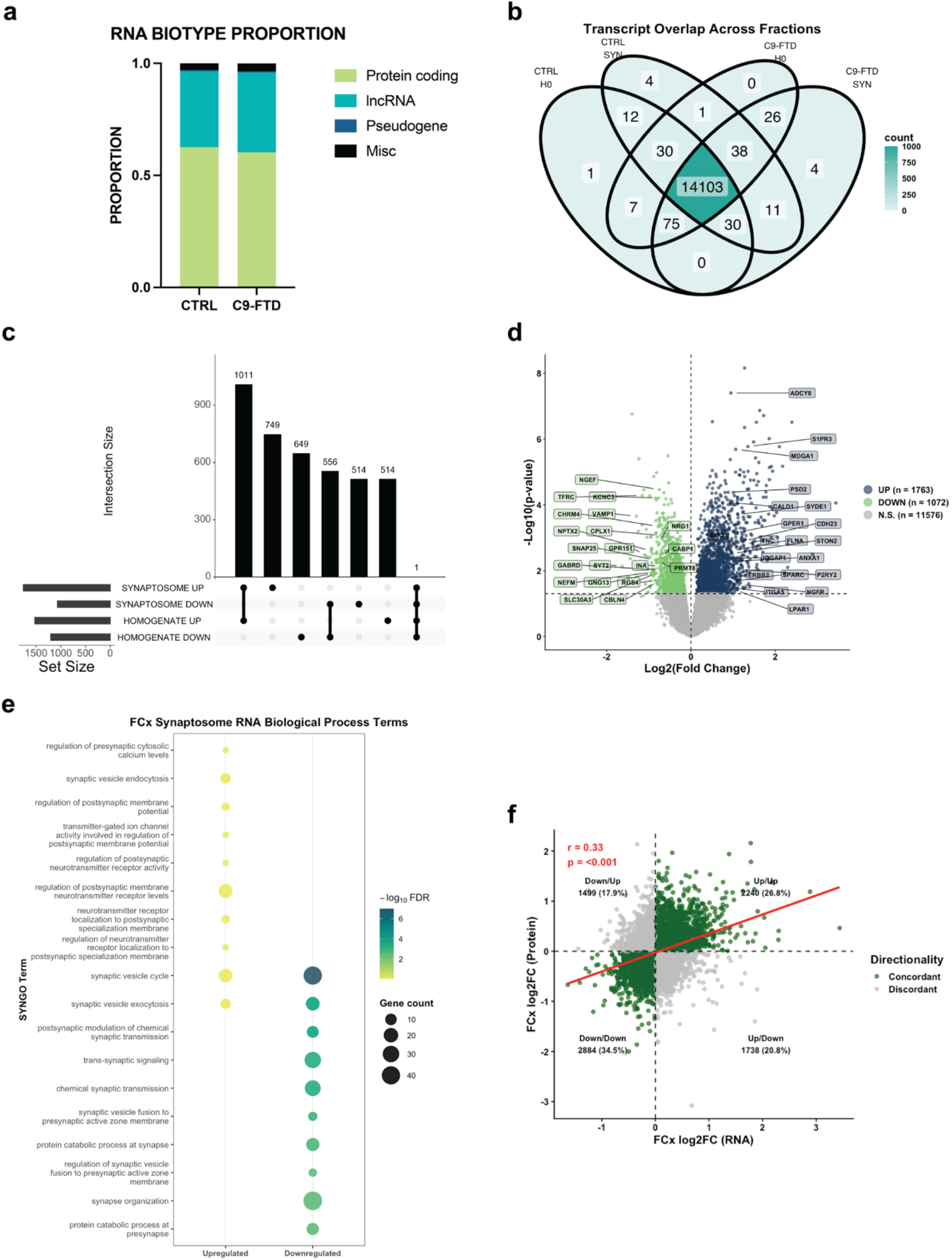
Frontal cortex-derived synaptosomal RNA reveals disease-associated aberrations and concordance with proteomic alterations. **a: RNA Biotype Composition:** Distribution of RNA biotypes (protein-coding, long non-coding RNA, and pseudogenes) across control and C9-FTD synaptosome fractions. **b: Transcript overlap across fractions:** Venn diagram of protein-coding RNA detected across H0 and P2 fractions in control and C9-FTD, defined as genes with >10 normalized counts in >50% of samples. **c: Compartment-specific differentially expressed genes:** UpSet plot displaying the quantity and overlap of significantly differentially expressed genes (DEG) across H0 and P2 fractions for protein-coding RNA, stratified by directionality (upregulated, log_2_FC≥0; downregulated, log_2_FC≤0). **d: Synaptosome Differential Expression:** Volcano plot of DEG identified in C9-FTD synaptosomes (n=2,835), highlighting the top 20 synaptic RNAs annotated based on absolute log_2_FC and SYNGO annotation. A total of 1,763 genes were significantly upregulated, and 1,072 genes were downregulated (DESeq2 Wald test, Benjamini-Hochberg adjusted p<0.05). Y axis represents -Log10(p-value). **e: Bi-directional Pathway Dysregulation:** SYNGO Biological Process analysis. While certain pathways show polarized regulation (e.g., calcium signaling is predominantly upregulated), core synaptic functions including **synaptic vesicle cycle** and **chemical synaptic transmission** exhibit significant alterations in both directions. SYNGO Biological Process enrichment analysis identified alterations in multiple presynaptic and postsynaptic pathways, with several core synaptic functions represented among both upregulated and downregulated genes. **f: Concordance between transcriptomic and proteomic alterations:** Scatter plot showing the correlation between synaptosomal RNA and protein log_2_FC. The “ball-like” distribution and moderate correlation coefficient (Pearson *r*=*0.33*) highlight a significant divergence between transcription and translation. The **concordant quadrants** (dark green) identify a subset of genes where RNA changes successfully translate to the proteome. In contrast, the **discordant quadrants** (grey) reveal a widespread indicating only partial concordance between transcriptomic and proteomic alterations.

To assess transcript distribution and localization across subcellular fractions, we compared protein-coding transcripts detected in homogenate and synaptosome fractions, focusing on protein-coding genes to facilitate integration with the corresponding proteomic dataset. Genes with >10 normalized counts in more than 50% of samples were considered detected. While substantial overlap was observed between fractions, distinct populations of compartment-associated transcripts were also observed (Fig.3b).

Differential expression analyses revealed widespread transcriptomic changes within C9-FTD synaptosomes. Overlap analysis demonstrated extensive compartment-associated differential gene expression in protein-coding transcripts across homogenate and synaptosome fractions, with a predominance of upregulated transcripts detected in synaptosomes (Fig.3c). We identified 2,835 differentially expressed genes (DEGs) in C9-FTD including 1,763 upregulated and 1,072 downregulated transcripts relative to controls (DESeq2 Wald test, p<0.05; Fig.3d).

To define the functional pathways most affected, SYNGO BP enrichment analyses were performed on upregulated and downregulated genes separately. Similar to the proteomic datasets, upregulated pathways showed modest enrichments for processes related to postsynaptic regulation and modulation of receptor signaling, including SYNGO BP terms: *regulation of postsynaptic membrane neurotransmitter receptor levels; regulation of postsynaptic neurotransmitter receptor activity; regulation of neurotransmitter receptor localization to postsynaptic specialization membrane; regulation of postsynaptic membrane potential; transmitter-gated ion channel activity involved in regulation of postsynaptic membrane potential; regulation of presynaptic cytosolic calcium levels; synaptic vesicle endocytosis*. In contrast, downregulated genes showed stronger enrichments in terms related to synaptic vesicle cycling and trans-synaptic communication, including SYNGO BP terms: *synaptic vesicle cycle; synaptic vesicle exocytosis; synaptic vesicle fusion to presynaptic active zone membrane; regulation of synaptic vesicle fusion to presynaptic active zone membrane; chemical synaptic transmission; trans-synaptic signaling; postsynaptic modulation of chemical synaptic transmission; synapse organization; protein catabolic process at synapse; protein catabolic process at presynapse*. Similarly, shared terms between upregulated and downregulated terms include synaptic vesicle cycle and synaptic vesicle exocytosis (Fig.3e). Curiously, similar to the synaptosomal proteomic alterations in C9-FTD frontal cortex, transcriptomic datasets revealed fewer enriched upregulated biological processes, whereas downregulated biological processes exhibited a more significant enrichment (based on -log10(False Discovery Rate-corrected p-value).

Given the extensive transcriptomic alterations identified within synaptosomes, we next examined the relationship between RNA and protein changes in C9-FTD synapses. Comparative analyses of RNA and protein log_2_ fold changes demonstrated only a moderate correlation between RNA and protein abundance (Pearson correlation r=0.33, p<0.001; Fig.3f). Although a subset of genes exhibited concordant directional changes across both datasets (61.3%), a subset (38.7%) displayed discordant RNA-protein relationships, suggesting impaired post-transcriptional regulation and molecular decoupling within diseased synapses.

### C9-FTD iPSC-cortical neurons display structural and functional synaptic deficits *in vitro*

To complement our analyses of human frontal cortex tissue and determine whether key synaptic features of C9-FTD could be recapitulated in vitro, patient-derived induced pluripotent stem cells (iPSCs) were differentiated into forebrain-like excitatory cortical neurons using dual-SMAD inhibition [52] and matured in monoculture to day in vitro (DIV) 100 prior to downstream analyses (Fig. 4a). Longitudinal Multielectrode Array (MEA) recordings were obtained between DiV65 and DiV100 to assess neuronal network activity, whereas immunofluorescence and synaptosome isolation experiments were conducted only at DiV100.

To determine whether C9-FTD cortical neurons exhibit structural synapse abnormalities, excitatory synapses were assessed by immunofluorescent labeling of presynaptic marker VGLUT1 and postsynaptic marker Homer1 (Fig.4b). Quantification of VGLUT1 puncta density normalized to MAP2-positive neurite volume revealed no difference between control and C9-FTD cortical neurons (p=0.6822; Mann-Whitney test), suggesting preservation of excitatory presynaptic terminals (Fig.4c). In contrast, Homer1 puncta density was reduced in C9-FTD cortical neurons (p<0.05; Mann-Whitney test), indicating selective vulnerability to excitatory postsynaptic structures (Fig.4d). Despite this postsynaptic reduction, the density of colocalized VGLUT1-Homer1 puncta was unchanged in C9-FTD (p=0.49; Mann-Whitney test), indicating that postsynaptic marker reductions occurred without impacting overall excitatory synapse density (Fig.4e). We next assessed inhibitory synapses using immunofluorescent labeling the inhibitory presynaptic marker VGAT and postsynaptic marker Gephyrin (Fig.4f). Similar to excitatory synapses, VGAT puncta density was unchanged between groups (p=0.468; Mann-Whitney test), indicating the preservation of inhibitory presynaptic terminals (Fig.4g). In contrast, Gephyrin puncta density was significantly reduced in C9-FTD cortical neurons (p<0.05; Mann-Whitney test), consistent with postsynaptic vulnerability at inhibitory synapses (Fig.4h). Unlike excitatory synapses, quantification of colocalized VGAT-Gephyrin puncta demonstrated a reduction in overall inhibitory synapse density in C9-FTD cortical neurons (p<0.05; Mann-Whitney test), suggesting a selective impairment of inhibitory synaptic connectivity (Fig.4i).

**Figure 4.**
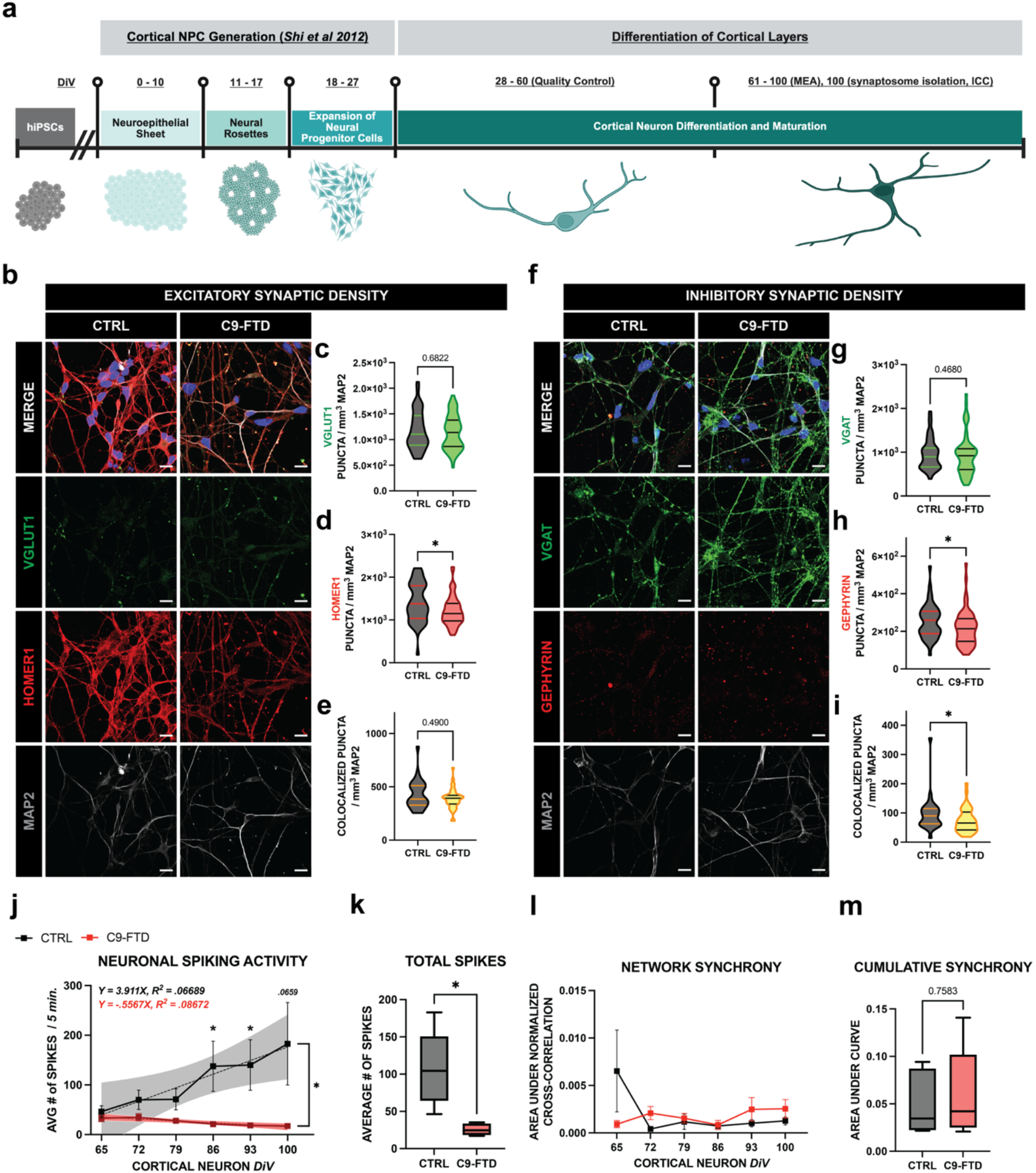
Structural and Functional Synaptic Deficits in C9-FTD iPSC-Derived Cortical Neurons. **a: iPSC Differentiation and Experimental Timeline:** Schematic diagram illustrating the differentiation of patient-derived iPSC-cortical neurons following a dual-SMAD inhibition protocol. iPSCs are differentiated into neuroepithelial sheets and neural rosettes, expanded as neural progenitor cells (NPC), and subsequently matured into excitatory, cortical neurons. Multielectrode Array (MEA) recordings were performed from DiV65 to DiV100, while synaptosome isolations and immunocytochemistry (ICC) experiments were performed at DiV100. **b-e: Excitatory Synapse Quantification:** Immunocytochemical staining of DiV100 cortical neurons for excitatory synaptic markers assessed by immunofluorescence for presynaptic VGLUT1 and postsynaptic Homer1 puncta. *Scale bar=10μM.* **c:** Quantification of VGLUT1 puncta density in DiV100 cortical neurons. Puncta density normalized to neuronal MAP2 volume (p=.6822; Mann-Whitney test). **d:** Quantification of HOMER1 puncta density in DiV100 cortical neurons. Puncta density normalized to neuronal MAP2 volume (*p<0.05; Mann-Whitney test), showing reductions in C9-FTD. **e:** Quantification of colocalized VGLUT1 and HOMER1 excitatory puncta density in DiV100 cortical neurons. Puncta density normalized to neuronal MAP2 volume (p=.49; Mann-Whitney test). **f: Inhibitory Synapse Quantification:** Immunocytochemical staining of DiV100 cortical neurons for inhibitory synaptic markers assessed by immunofluorescence for presynaptic VGAT and postsynaptic Gephyrin puncta. *Scale bar=10μM.* **g:** Quantification of VGAT puncta density in DiV100 cortical neurons. Puncta density normalized to neuronal MAP2 volume (p=.4680; Mann-Whitney test). **h:** Quantification of Gephyrin puncta density in DiV100 cortical neurons. Puncta density normalized to neuronal MAP2 volume (*p<0.05; Mann-Whitney test), showing reductions in C9-FTD. **i:** Quantification of colocalized VGAT and Gephyrin inhibitory puncta density in DiV100 cortical neurons. Puncta density normalized to neuronal MAP2 volume (p<0.05; Mann-Whitney test), showing reductions in colocalized inhibitory synapses in C9-FTD. **j: Longitudinal Neuronal Spiking Activity:** Functional assessment of cortical neurons using Multielectrode Array (MEA) analyses, demonstrating a progressive decrease in network activity in C9-FTD iPSC-CN, as measured by the number of spikes over time (p<0.05, Welch’s t-test; p<0.0001, 2-way ANOVA). **k:** Box plot quantifying the total number of spikes across DiV65 to DiV100 in control and C9-FTD, showing decreased numbers of total spikes in C9-FTD (p<0.05; Welch’s t-test). **l: Network Synchrony:** Longitudinal assessment of network synchrony in control and C9-FTD cortical neurons, represented as area under normalized cross-correlation. **m:** Cumulative network synchrony quantified as the area under the normalized cross-correlation curve.DiV65 to DiV100 in control and C9-FTD cortical neurons (p=.7583; Welch’s t-test).

We wondered whether the structural abnormalities were accompanied by impaired neuronal network function; we performed longitudinal MEA recordings across cortical neuron maturation. C9-FTD cortical neurons exhibited a progressive reduction in the number of spontaneous neuronal spiking activity between DiV65 and DiV100 (p<0.05; 2-way ANOVA; Fig.4j). Quantification of cumulative spiking activity further confirmed reduced overall neuronal firing in C9-FTD cultures (p<0.05; Welch’s t-test) (Fig.4k), consistent with impaired network excitability. We next examined if disease-associated changes manifested in network-level synchronous events. In contrast to the spiking activity, longitudinal assessment of normalized cross-correlation revealed no differences in network synchrony across maturation (Fig.4l), and cumulative synchrony measurements were similarly unchanged between groups (p=0.7583; Welch’s t-test; Fig.4m), suggesting that synchronous network activity is preserved despite reduced neuronal firing.

Together, these findings indicate that C9-FTD iPSC-derived cortical neurons exhibit selective postsynaptic vulnerability, reduced inhibitory synapse density, and progressive hypoexcitability, supporting the presence of both structural and functional synaptic impairments in disease.

### iPSC-neuron derived synaptosomes recapitulate disease-associated pathways observed in C9-FTD frontal cortex

To determine whether patient-derived iPSC-cortical neurons recapitulate synaptic abnormalities observed in postmortem tissue, we performed proteomic analyses on homogenate and synaptosome fractions from control and C9-FTD neurons. We first verified the integrity and enrichment of the synaptosome fraction by western blotting for the presynaptic marker Synaptophysin (SYP) and postsynaptic density marker PSD-95 (Supplement Fig.2a). PCA of the iPSC proteomic dataset showed clear separation of homogenate and synaptosome fractions, with disease status explaining the largest proportion of variance (PC1=40.6%), followed by fraction identity (PC2=23.3%; Supplement Fig.2b). Consistent with these findings, comparison of compartment-specific protein abundance demonstrated enrichment of synaptic markers in P2 fractions and nuclear-associated markers in H0 fractions, confirming successful subcellular fractionation and synaptic enrichment (Supplement Fig.2c). PCA of the transcriptomic data similarly separated H0 and P2 fractions, again with disease status accounting for the greatest sources of variance (PC1=47.1%), and fraction identity contributing additional variance (PC2=16.4%; Supplement Fig.2d). In agreement with the proteomic dataset, P2 fractions were enriched for synaptic transcripts relative to H0 fractions (Supplement Fig.2e), supporting successful isolation of synapse-associated RNA.

Comparative analysis across biochemical fractions revealed both global and synapse-specific alterations in the *in vitro* system. Differential abundance analysis identified protein dysregulation across both homogenate and synaptosome fractions in C9-FTD relative to non-neurological disease controls. Overall, 14.6% of proteins detected in the homogenate fraction and 12.4% of proteins detected in the synaptosome fraction were altered in C9-FTD (p<0.05), indicating substantial proteomic alterations in both biochemical fractions (Fig.5a). Analysis of protein detection across fractions in > 66% of samples revealed substantial overlap between compartments, while also identifying synapse-specific signatures in control and C9-FTD synaptosomes (Fig.5b).

To define compartment-specific disease-associated changes, we quantified overlap among DAPs (p<0.05) across homogenate and synaptosome fractions, stratified by direction of change. Analysis of DAP overlap across homogenate and synaptosome fractions revealed a subset of proteins preferentially detected in the synaptosome fraction, suggesting that many disease-associated proteomic changes are synaptically enriched in an iPSC-cortical neuron model (Fig. 5c). We therefore focused subsequent analyses on the synaptosome fraction. Differential abundance analysis of the synaptic proteome identified 962 DAPs in C9-FTD, comprising 508 upregulated and 454 downregulated proteins relative to control (p<0.05; Fig.5d). Significant proteins identified in the synaptosome fraction (p<0.05) were ranked by log_2_ fold changes and then annotated using the SYNGO database. The highest-ranked synaptic proteins included proteins associated with vesicle trafficking and synapse organization, suggesting remodeling of both presynaptic and postsynaptic compartments in C9-FTD.

To investigate the functional consequences most strongly affected in C9-FTD iPSC-cortical neuron synapses, we performed SYNGO BP enrichment analysis on synaptosomal DEPs, separately considering upregulated and downregulated proteins. Upregulated proteins showed modest enrichment in pathways related to structural remodeling and postsynaptic organization, including SYNGO BP terms: *postsynapse organization; regulation of postsynapse organization; synapse adhesion between pre- and post-synapse; translation at presynapse; translation at postsynapse*. Conversely, downregulated proteins were more strongly enriched for pathways related to synaptic signaling and cytoskeletal architecture, with SYNGO BP terms: *chemical synaptic transmission; postsynaptic cytoskeleton organization; postsynaptic actin cytoskeleton organization; synaptic target recognition; maintenance of synapse structure*. Several overrepresented terms related to synapse assembly and organization were shared between upregulated and downregulated gene sets, including *synapse organization, synapse assembly, trans-synaptic signaling, regulation of synapse assembly, and regulation of postsynaptic membrane neurotransmitter receptor levels*, with 50% of terms overlapping in both upregulated and downregulated directions (Fig.5e). Interestingly, multiple enriched pathways were also observed in C9-FTD postmortem frontal cortex tissue, indicating that patient-derived iPSC-cortical neurons recapitulate key features of C9-FTD pathology *in vitro*.

**Figure 5.**
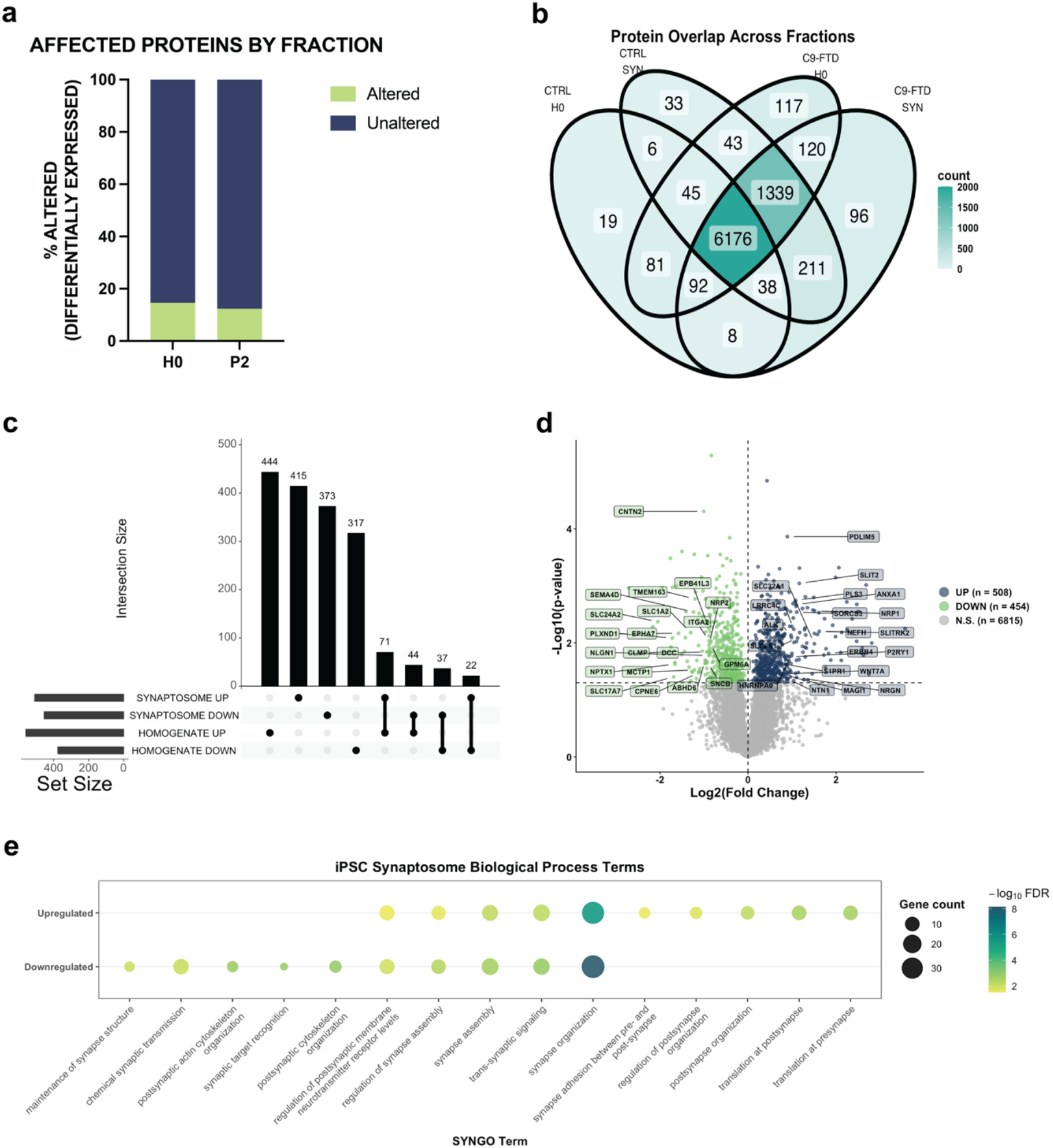
iPSC-derived synaptosomes recapitulate disease-associated features and pathways of C9-FTD. **a: Protein Alteration by Fraction:** Proportion of significantly altered protein expression (differential expression, p=<0.05) identified in H0 and P2 fractions following differential proteomic analysis between control and C9-FTD cases in frontal cortex tissue (14.6% H0, 12.4% P2; alterations relative to C9-FTD). **b: Proteomic Overlap:** Venn diagram displaying the distribution of proteins identified across H0 and P2 fractions in control and C9-FTD iPSC cell lines. **c: Intersection of Differentially Expressed Proteins (DEPs):** UpSet plot displaying the quantity and overlap of significantly differentially expressed proteins (DEP) across H0 and P2 fractions, stratified by directionality (upregulated, log_2_FC≥0; downregulated, log_2_FC≤0). **d: iPSC Synaptosome Differential Expression:** Volcano plot of DEP identified in C9-FTD synaptosomes (n=962), highlighting the top 20 synaptic genes annotated based on absolute log_2_FC and SYNGO annotation. A total of 508 proteins were significantly upregulated, and 454 proteins were downregulated (p=<0.05, *t-test*). **e: Functional Pathway Recapitulation:** SYNGO Biological Process enrichment analysis of significantly upregulated and downregulated proteins in synaptosome fractions, displaying the top 10 enriched unique and shared terms per direction (5 unique and 5 shared terms) (p=<0.05, relative to C9-FTD).

### Synaptosomal transcriptomics reveal transcriptional dysregulation and RNA-protein concordance in C9-FTD iPSC-cortical neurons

To determine whether the transcriptomic abnormalities observed in C9-FTD postmortem synaptosomes are recapitulated in patient-derived iPSC-cortical neurons, we compared the transcriptomes of homogenate and synaptosome fractions. Analysis of RNA biotype composition showed that protein-coding transcripts comprised the predominant RNA species in both control and C9-FTD samples (>80%), at a higher proportion than in postmortem tissue, with additional contributions from lncRNA (Fig.6a). RNA biotype composition was comparable between control and C9-FTD groups, indicating preservation of transcriptome organization in the iPSC-cortical neuron model.

To assess transcript distribution across compartments, protein-coding genes were considered detected when they showed >10 normalized counts in >66% of samples. Comparative analysis revealed substantial overlap in transcript detection between homogenate and synaptosome fractions, while also identifying distinct populations of compartment-specific transcripts (Fig. 6b). Differential expression analysis identified transcriptional aberrations in C9-FTD iPSC-cortical neurons. Overlap analysis demonstrated both shared and synapse-specific differential gene expression across homogenate and synaptosome fractions, stratified by directionality (Fig.6c). In contrast to the proteomic data, the largest overlapping groups consisted of protein-coding genes that were downregulated in both fractions, suggesting an overall transcriptional suppression in the iPSC model. To focus on synaptic transcriptomic alterations, subsequent analyses were restricted to protein-coding genes detected in the synaptosome fractions. This analysis identified 402 protein-coding DEGs in C9-FTD iPSC-derived synaptosomes, including 138 upregulated and 264 downregulated genes relative to controls (DESeq2 Wald test, p<0.05; Fig.6d). Annotation using the SYNGO database together with selection of genes showing the largest log_2_ fold changes, revealed widespread dysregulation of genes involved in synaptic vesicle trafficking and postsynaptic organization, suggesting altered expression of genes governing neurotransmission and synaptic architecture in C9-FTD.

**Figure 6.**
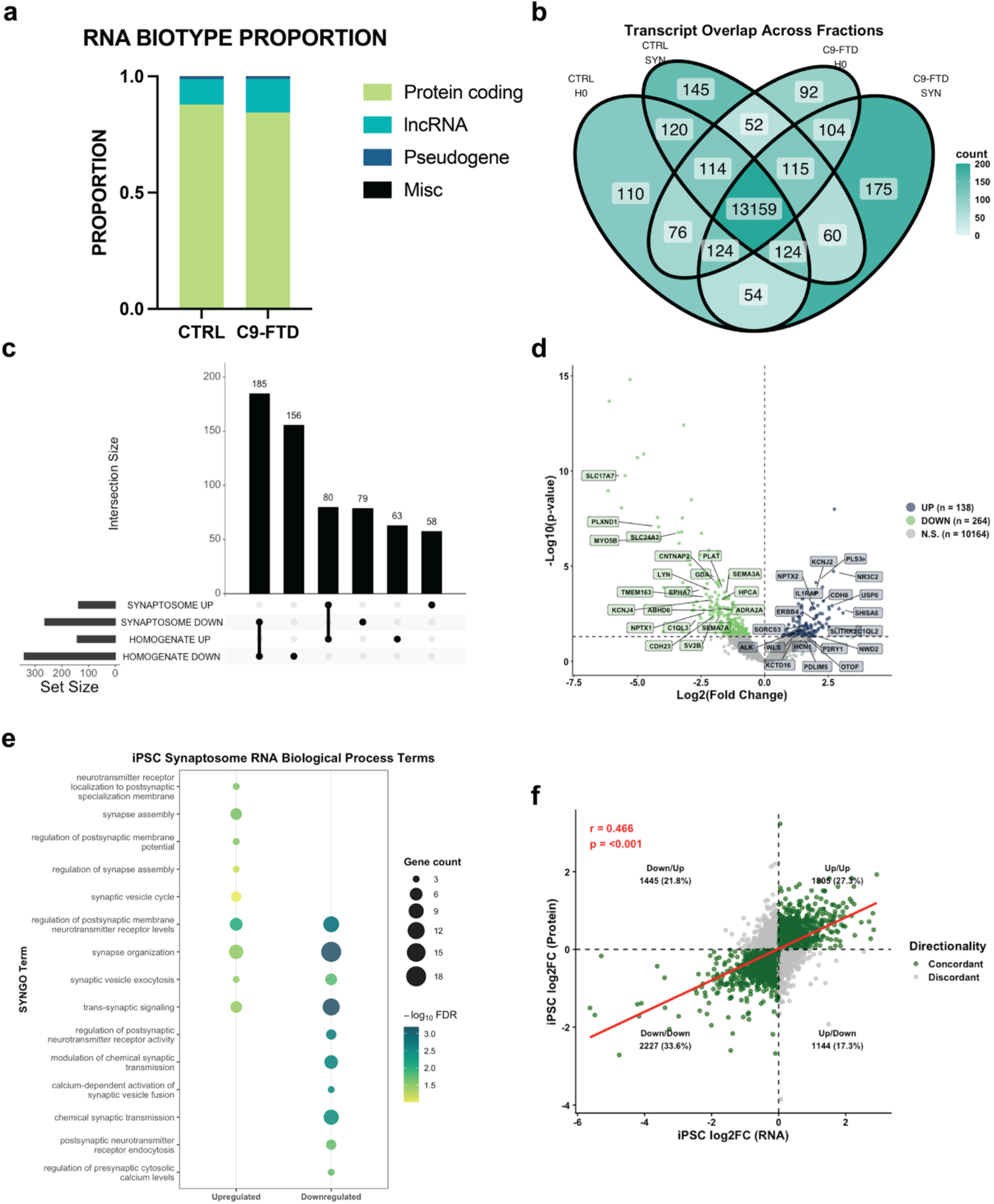
iPSC-derived synaptosomal RNA reveals disease-associated aberrations and concordance with proteomic alterations. **a: RNA Biotype Proportions:** Distribution of RNA biotypes (protein-coding, long non-coding RNA, and pseudogenes) across control and C9-FTD synaptosome fractions. **b: Transcriptional Overlap and Compartmentalization:** Venn diagram of protein-coding RNA detected across H0 and P2 fractions in control and C9-FTD, defined as genes with >10 normalized counts in >66% of samples. **c: Transcriptional Overlap and Compartmentalization:** The largest intersection consisted of genes that were downregulated in both homogenate and synaptosome fractions, indicating that many transcriptional changes are shared across compartments. **d: Synaptosome-Specific DEGs:** Volcano plot of protein-coding DEG identified in C9-FTD synaptosomes (n=402), highlighting the top 20 synaptic genes annotated based on absolute log_2_FC and SYNGO annotation. A total of 138 genes were significantly upregulated, and 264 genes were downregulated (DESeq2 Wald test, p<0.05). Y axis represents -Log10(p-value). **e: Pathway Enrichment:** SYNGO Biological Process enrichment analysis of significantly upregulated and downregulated protein-coding RNA in synaptosome fractions showing the top enriched SYNGO Biological Process terms among upregulated and downregulated genes. **f: Enhanced RNA-Protein Concordance in iPSCs:** Correlation analysis of synaptosomal transcriptome (x-axis) and proteome (y-axis) log_2_FC. The correlation (Pearson r=0.466, p < 0.001) is higher than that observed in post-mortem tissue, though the presence of discordant quadrants (grey) is still consistent with significant post-transcriptional regulation. The high percentage of concordant downregulation (33.6%) is consistent with reduced protein levels at the iPSC synapse are partly driven by decreased transcript availability.

To define the functional consequences of these transcriptomic changes, we conducted Biological Process enrichment analysis on synaptosomal DEGs using the SYNGO database. As observed in the iPSC proteomics dataset, upregulated gene enrichment was modest and reflected processes involved in synaptic structural organization and vesicle dynamics, including SYNGO BP terms: *synaptic vesicle cycle; synapse assembly; regulation of synapse assembly; regulation of postsynaptic membrane potential; neurotransmitter receptor localization to postsynaptic specialization membrane*. In contrast, downregulated genes showed stronger enrichment for pathways related to neurotransmitter release modulation, including SYNGO BP terms: *chemical synaptic transmission; modulation of chemical synaptic transmission; regulation of postsynaptic neurotransmitter receptor activity; postsynaptic neurotransmitter receptor endocytosis; regulation of presynaptic cytosolic calcium levels; calcium-dependent activation of synaptic vesicle fusion*. Several pathways were shared between upregulated and downregulated gene sets, including *synapse organization, synaptic vesicle exocytosis, trans-synaptic signaling, and regulation of postsynaptic membrane neurotransmitter receptor levels* (Fig.6e). Consistent with iPSC synaptosome proteomics, transcriptomic analyses showed modest upregulation and increased enrichment of downregulated terms relative to C9-FTD. Several enriched pathways overlapped with those identified in postmortem transcriptional profiles, supporting the conclusion that patient-derived iPSC-cortical neurons recapitulate key features of disease-associated synapse-specific transcriptomic disruption observed in C9-FTD brain tissue.

Given the concurrent alterations in protein abundance and RNA expression, we next assessed the relationship between proteomic and transcriptomic alterations in iPSC-derived synaptosomes. Correlation analysis of RNA and protein log_2_FC revealed a moderate positive correlation between transcript and protein abundance changes (Pearson r = 0.466, p < 0.001; Fig. 6f), exceeding the concordance observed in frontal cortex synaptosomes. Most genes exhibited concordant changes at the transcript and proteins levels (>60%), whereas the remaining genes displayed discordant RNA-protein coupling. The predominance of concordant downregulation indicates that decreases in synaptic protein abundance are frequently accompanied by corresponding reductions in transcript abundance in the iPSC-derived cortical neuron model. Together, these findings demonstrate that iPSC-derived synaptosomes recapitulate key molecular features of C9-FTD synaptic dysfunction through both transcriptional and post-transcriptional regulation

### Cross-model multi-omic integration identifies conserved pathways of synaptic dysfunction in C9-FTD

To identify synaptic molecular alterations shared between postmortem tissue and the iPSC model, we performed integrated cross-model analyses of the proteomic and transcriptomic datasets from both systems. Comparative integration of the four datasets allowed us to identify conserved molecular alterations and shared biological process pathways associated with synaptic dysfunction in C9-FTD. Overlap analysis of DEGs and DAPs identified in synaptosomes from frontal cortex and iPSC-cortical neurons revealed both model-specific and inter-model alterations across protein and RNA datasets (Fig.7a Although many alterations were unique to individual datasets, several altered genes and proteins were consistently identified in at least three of the four datasets, highlighting molecular changes that are reproducible across independent experimental systems and therefore represent high-confidence candidates for C9-FTD-associated synaptic dysfunction.

To determine whether these shared molecular changes converged on common biological pathways, SYNGO Biological Process enrichment analysis was performed separately on overlapping DAPs and DEGs identified across frontal cortex and iPSC-derived datasets (Fig.7b). This analysis revealed shared synaptic biological processes associated with synapse assembly, organization, and signaling, particularly within the proteomic datasets, which demonstrated a greater degree of overlap. Conversely, transcriptomic datasets showed less overlap overall, converging primarily on trans-synaptic signaling pathways. These finding suggest that while transcriptional aberrations may be more heterogenous across model systems, they ultimately converge on shared synaptic processes that are reflected more consistently at the protein level.

To further visualize conserved and divergent molecular signatures across datasets, integrated multi-omic heatmaps were generated using the top 25 genes associated with the highest enriched SYNGO Biological Process terms, ranked by the summed absolute log_2_FC values (Fig.7c-g). Comparative analysis demonstrated convergent patterns of dysregulation across protein and RNA datasets associated with *synapse organization* (Fig.7c), *regulation of synapse assembly (*Fig.7d), *regulation of postsynaptic membrane neurotransmitter receptor levels* (Fig.7e), *synapse assembly* (Fig.7f), and *trans-synaptic signaling* (Fig.7g). While the magnitude of abundance or expression changes varied across datasets and model systems, the majority of genes retained consistent directionality, supporting the presence of conserved pathway-level abnormalities underlying C9-FTD synaptic dysfunction.

Collectively, these findings indicate that patient-derived iPSC cortical neurons reproduce key molecular signatures identified in postmortem C9-FTD synaptosomes, highlighting conserved pathways involved in synapse assembly and synapse organization across independent experimental systems.

To further quantify the degree of molecular overlap between model systems, correlation analyses comparing log_2_FC values across datasets were performed. Protein-level comparisons between frontal cortex and iPSC-derived synaptosomes demonstrated only modest overall concordance (Pearson correlation r=0.044, p<0.001; Supplement Fig.3a). Similarly, transcriptomic comparisons revealed minimal correlation between model systems (Pearson correlation r=0.074, p<0.001; Supplement Fig.3b). Despite limited concordance at the individual gene and protein level, both model systems converged on shared synaptic pathways, indicating that iPSC-cortical neuron-derived synaptosomes preserve core biological processes underlying C9-FTD-associated synaptic dysfunction, while only partially recapitulating the broader molecular landscape present in postmortem frontal cortex synaptosomes.

### Cryptic exon analysis reveals synaptic localization of TDP-43-associated cryptic exon-containing transcripts in C9-FTD

Given confirmed FTLD-TDP A pathology in these C9-FTD frontal cortex cases, we next investigated whether transcripts containing TDP-43-associated CE are present in synaptosome fractions. We first assessed CE inclusion in *STMN2*, a canonical marker of TDP-43 loss-of-function splicing dysregulation. Visualization of synaptosome-derived RNA using Integrated Genome Viewer (IGV) sashimi plots revealed aberrant CE-containing splice junctions in C9-FTD cases that were absent in control (Fig.8a). Quantification using percent spliced in (PSI) quantification confirmed increased inclusion of the STMN2 cryptic exon in C9-FTD synaptosomes (p=0.0007; Mann-Whitney test; Fig.8b). We next examined *KALRN*, a key synaptic scaffolding and signaling gene previously implicated in TDP-43 associated splicing alterations [18, 23]. Sashimi plot analyses demonstrated the presence of *KALRN* CE-containing transcripts in C9-FTD synaptosomes (Fig.8c). However, PSI quantification revealed greater inter-individual variability and low-level CE inclusion in several control cases, resulting in no significant group difference (p=0.1462; Mann-Whitney test; Fig.8d). In contrast, analysis of *UNC13A*, a key regulator of synaptic vesicle release and established TDP-43 splicing substrate, showed no detectable CE-containing transcripts within synaptosome fractions from either control or C9-FTD cases (p>0.9999; Mann-Whitney test; Fig.8e-f). These findings suggest selective synaptic localization of specific CE-containing transcripts.

To determine whether synaptic CE accumulation was associated with altered synaptic transcript abundance, we next assessed normalized gene expression of the canonical transcripts within the synaptosome. Expression of *STMN2*, *KALRN*, and *UNC13A* mRNA showed no changes between control and C9-FTD case (Fig.8g-i), suggesting that the presence of local splicing aberrations was not simply driven by altered transcript abundance. Similarly, synaptosomal protein analyses revealed no changes in *STMN2* or *KALRN* (Fig.8j-k) proteins levels, whereas UNC13A protein expression showed a trend toward reduction in C9-FTD cases (p=0.0502; Fig.8i). Together, these findings suggest that local CE-associated splicing aberrations occur independently of major changes in steady-state synaptic RNA or protein abundance.

To determine whether these observations are reflected within the frontal cortex homogenate fraction, we expanded our investigation of CE-containing transcripts to whole brain tissue lysates (Supplement Fig.4a-f). CE-containing transcripts in *STMN2* and *KALRN* were absent in control homogenates, while minimal presence of *KALRN* events were detected in the synaptosome (Table 3). In contrast to synaptosomes, C9-FTD homogenates demonstrated robust CE inclusion across all investigated genes, *STMN2, KALRN, UNC13A*, consistent with widespread TDP-43 splicing dysfunction in bulk tissue containing nuclear-localized transcripts (Supplement Fig.4a,c,e). PSI analyses confirmed increased CE inclusions for *STMN2* (p<0.0001); *KALRN* (p=0.0031); and *UNC13A* (p=0.0108) in C9-FTD homogenates (Mann-Whitney test; Supplement Fig.4b,d,f).

**Table 3.**
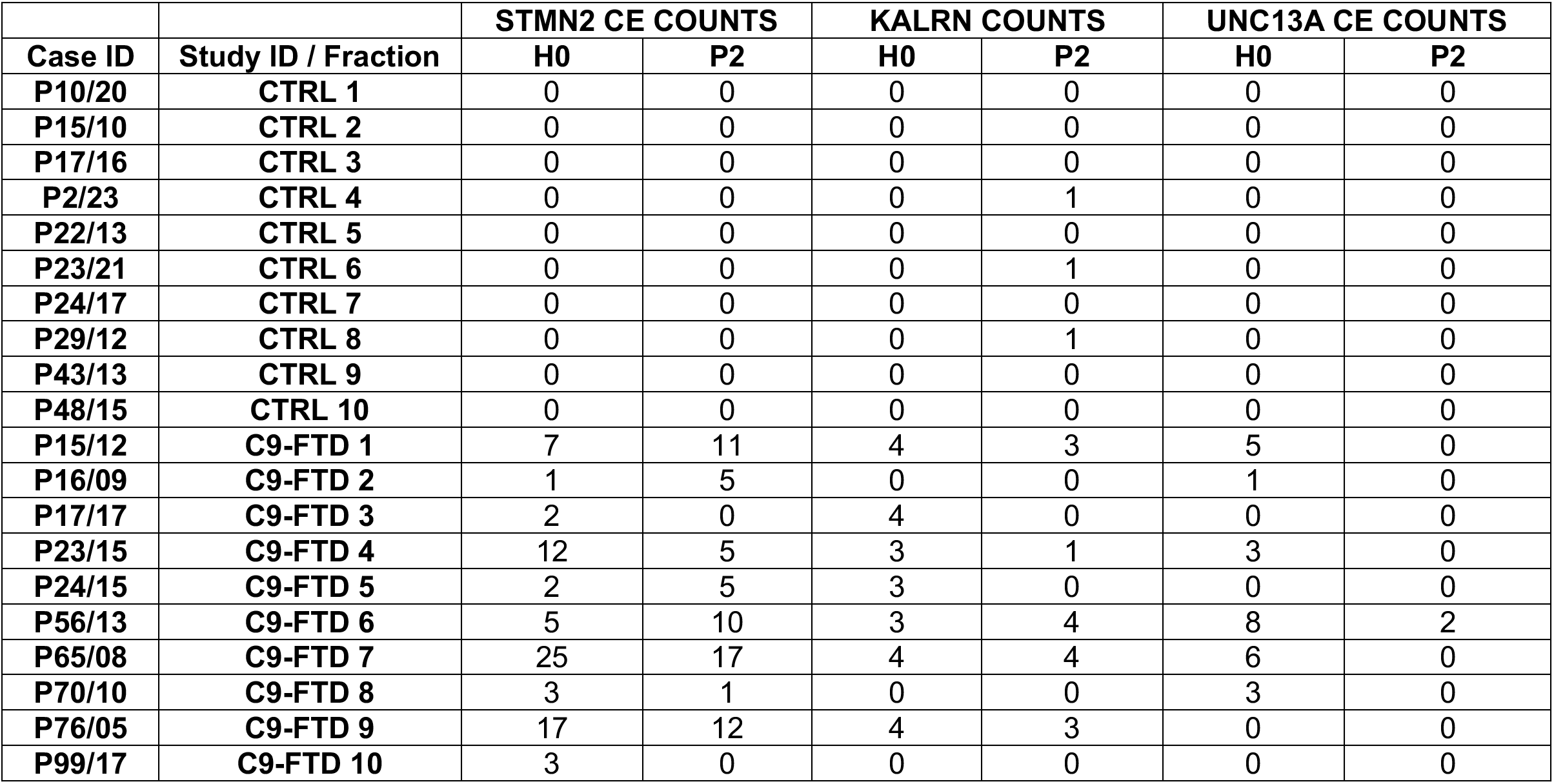
Detection of total Cryptic Exon (CE) counts for *STMN2, KALRN, UNC13A* across frontal cortex case condition and fraction.

To further assess subcellular localization patterns, synaptosome to homogenate abundance ratios (P2:H0) were calculated. *STMN2* exhibited the highest relative synaptic enrichment (P2:H0=0.8571), followed by *KALRN* (P2:H0=0.6), whereas *UNC13A* CE-containing transcripts showed minimal synaptic localization (P2:H0=0.0769; Table 4).

**Table 4.**
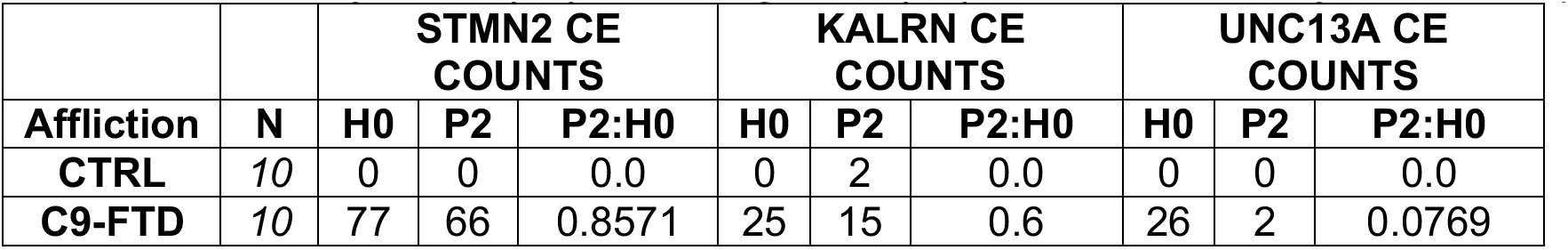
Ratio of synaptic (P2) to homogenate (H0) localization of Cryptic Exons (CE).

| Affliction | N | STMN2 CE COUNTS |  |  | KALRN CE COUNTS |  |  | UNC13A CE COUNTS |  |  |
| --- | --- | --- | --- | --- | --- | --- | --- | --- | --- | --- |
|  |  | H0 | P2 | P2:H0 | H0 | P2 | P2:H0 | H0 | P2 | P2:H0 |
| CTRL | 10 | 0 | 0 | 0.0 | 0 | 2 | 0.0 | 0 | 0 | 0.0 |
| C9-FTD | 10 | 77 | 66 | 0.8571 | 25 | 15 | 0.6 | 26 | 2 | 0.0769 |

Collectively, these findings demonstrate that TDP-43-associated CE-containing transcripts are present within synaptic compartments in C9-FTD, identifying their localization to the synapse as a previously undescribed feature of TDP-43 proteinopathy.

## DISCUSSION

In this study, integrated proteomic and transcriptomic profiling of isolated synaptosomes from C9-FTD frontal cortex revealed extensive remodeling of synaptic molecular networks, convergent alterations in synapse organization and neurotransmission pathways, and the unexpected presence of TDP-43-associated cryptic exon-containing transcripts within synaptic compartments.

Differential abundance and expression analyses across C9-FTD and non-neurological disease control frontal cortex tissue samples showed widespread aberrations in synapse-specific pathways at both the protein (Fig.2) and RNA (Fig.3) level. Using SYNGO, we identified protein disruptions associated with synaptic assembly, organization, and modulation, and synaptic vesicle trafficking, with greater magnitude changes associated with downregulated processes. Transcriptomic profiling of synaptic-localized transcripts showed changes associated with energetic and ionic regulation of synaptic vesicle dynamics, exocytosis, and synaptic organization, further highlighting specific aspects of synaptic dysfunction. Our findings are consistent with prior work profiling synaptic proteomes in C9-ALS across premotor dorsolateral prefrontal cortex (Brodmann’s Area 9) and motor cortex (Brodmann’s Area 4) regions [34]. While this study provided insight into the spatial distribution across disease-affected regions in ALS and ALS with cognitive impairment, the study was limited to phenotypic ALS and proteomics analysis alone. Previous studies profiling the transcriptome of C9-ALS and C9-FTD tissues have primarily focused on bulk and single nuclei RNA sequencing, thus providing limited insight into localized synaptic RNA [18, 35]. By profiling both the synaptic proteome and local transcriptome in postmortem frontal cortex tissue (Fig.2,3) and patient-derived iPSC-cortical neurons (Fig.5,6), our study enabled a multi-omic and multi-model comparison across disease stage (Fig.7). Together, this provides conserved and disease progression specific mechanisms of C9-linked synaptic dysfunction and neurodegeneration.

**Figure 7.**
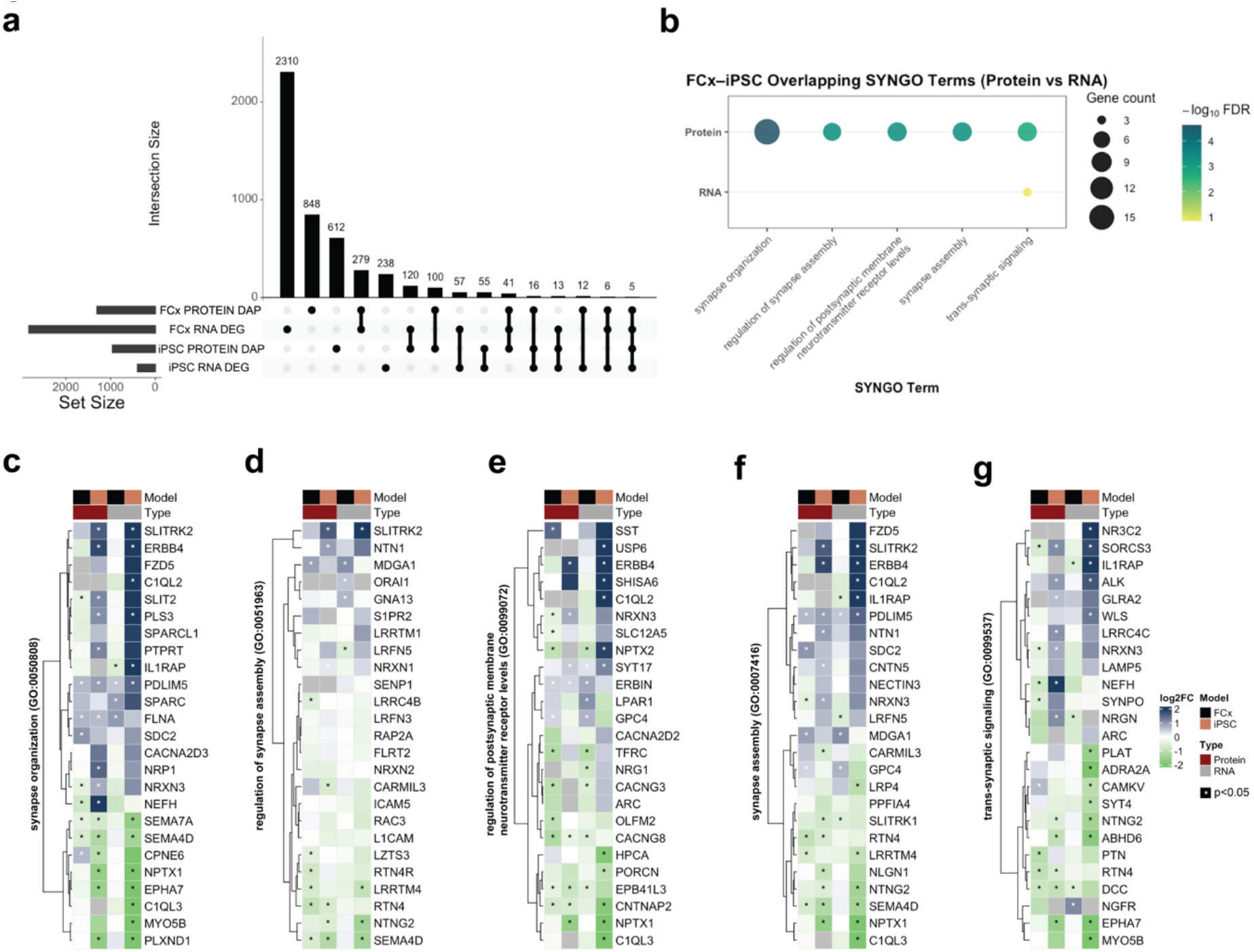
Cross-Model Integration Identifies Conserved Molecular Signatures of C9-FTD Synaptic Pathology. **a: Global Intersection of Brain and iPSC Data:** UpSet plot quantifying the overlap of differentially expressed genes (DEG) and abundant proteins (DAP) across FCx tissue and iPSC-derived synaptosomes. The presence of shared hits across all four datasets (right-most bars) identifies a subset of genes and proteins consistently altered across multiple datasets, highlighting conserved molecular signatures of C9-FTD synaptic pathology. **b: Conserved Biological Pathways:** SYNGO Biological Process enrichment analysis of the DEGs/DAPs shared between FCx and iPSC models. Key conserved processes include **synapse organization**, **synapse assembly**, and **trans-synaptic signaling**. **c–g: Integrated Signature Heatmaps:** heatmaps displaying log_2_FC values for the top 25 genes within conserved SYNGO terms: (**c**) synapse organization, (**d**) regulation of synapse assembly, (**e**) regulation of postsynaptic membrane neurotransmitter receptor levels, (**f**) synapse assembly, and (**g**) trans-synaptic signaling. These plots illustrate conserved expression patterns across human brain tissue and iPSC models for both RNA and protein modalities.

Functional analyses of network activity revealed decreased neuronal spikes in C9-FTD iPSC-cortical neurons, suggesting a phenotype of progressive hypoexcitability (Fig.4). These observations are in contrast with established models that demonstrate how the C9-HRE results in hyperexcitability from iPSC and mouse models [7, 42]. These discrepancies may represent differences associated with disease phenotype when comparing FTD to ALS and given the spatially distributed regions of selective vulnerability from frontal cortex to motor cortex and spinal cord in FTD and ALS, respectively, suggesting neuronal subtype vulnerability to synaptic dysfunction.

Both postmortem tissue and iPSC-cortical neuron-derived synaptosomes demonstrated a consistent pattern of decreased expression of key synaptic genes implicated in various neurodegenerative diseases, including *NPTX1, NPTX2,* and *NPTXR*. Neuronal Pentraxins and their receptors play a role in excitatory synapse formation [19], where the loss of these proteins have been associated with impaired synaptic transmission and reduced synaptic density [58, 59]. Recent CSF studies of genetic forms of FTD, including *C9ORF72, GRN,* and *MAPT* mutation carriers have shown reductions in *NPTX1* and *NPTXR* levels across patient cohorts [53], demonstrating *NPTX* dysregulation and synaptic degeneration in FTD. Our study demonstrates consistent downregulation of *NPTX1* and *NPTXR* across tissue and iPSC-derived synaptosomal proteomic and transcriptomic datasets. These results align with previous biomarker studies and further demonstrate synaptic vulnerability in C9-FTD, and that iPSC-cortical neuron models capture features of synapse loss observed in C9-FTD disease progression.

Aberrant CE inclusion resulting from nuclear depletion of TDP-43 is now recognized as a defining molecular feature of TDP-43 proteinopathies. CE-containing transcripts in TDP-43 targets, including *STMN2, UNC13A* and *KALRN*, have been extensively reported in ALS and FTD human postmortem tissue and experimental models, and are widely used as markers of TDP-43 loss-of-function [5, 31, 39]. Previous studies, including our single nucleus RNA sequencing analyses from C9-FTD cortex, have demonstrated robust CE inclusion within neuronal nuclei, supporting the concept that splicing dysregulation is a central consequence of TDP-43 pathology [18]. Recent evidence suggests that aberrantly spliced transcripts, including those affected by alternative polyadenylation events, are not necessarily retained within the nucleus and can be detected in cytoplasmic compartments, where they may contribute to disease-associated molecular dysfunction [6].

Building on these findings, our study provides evidence that CE-containing transcripts are present within isolated synaptosomal fractions from C9-FTD brain tissue, demonstrating that selected TDP-43-dependent splicing abnormalities extend into distal synaptic compartments (Fig.8). Among the transcripts examined, *STMN2* showed the strongest synaptic enrichment, *KALRN* displayed detectable but variable, case-dependent CE inclusion, whereas *UNC13A* CE-containing transcripts were largely absent from synaptosomes despite robust detection in matched homogenate fractions in C9-FTD. These findings argue against a generalized redistribution of all aberrantly spliced transcripts and instead suggest transcript-specific trafficking, retention, or stability within synaptic compartments. Given the importance of local RNA transport and translation for maintaining synaptic structure and function [24, 50], the presence of CE-containing transcripts at the synapse raises the possibility that TDP-43-dependent splicing abnormalities may directly contribute to synaptic dysfunction in C9-FTD and TDP-43 pathologies [23]. Notably, CE inclusion occurred in the absence of major changes in total transcript or protein abundance for *STMN2, KALRN,* or *UNC13A*, indicating that cryptic splicing can occur independently of steady-state gene expression changes. In the context of CE in *KALRN,* multiple isoforms of the gene exist (such as *KALRN*-7,-9,-12) [30], and while CE inclusions may affect a specific isoform, bulk RNA sequencing will not reflect this aberration. Together, these finding identify synaptic localization of CE-containing transcripts as a previously underappreciated feature of TDP-43 proteinopathy and suggest that disrupted local RNA homeostasis may represent an early molecular event contributing to synaptic vulnerability in C9-FTD.

**Figure 8.**
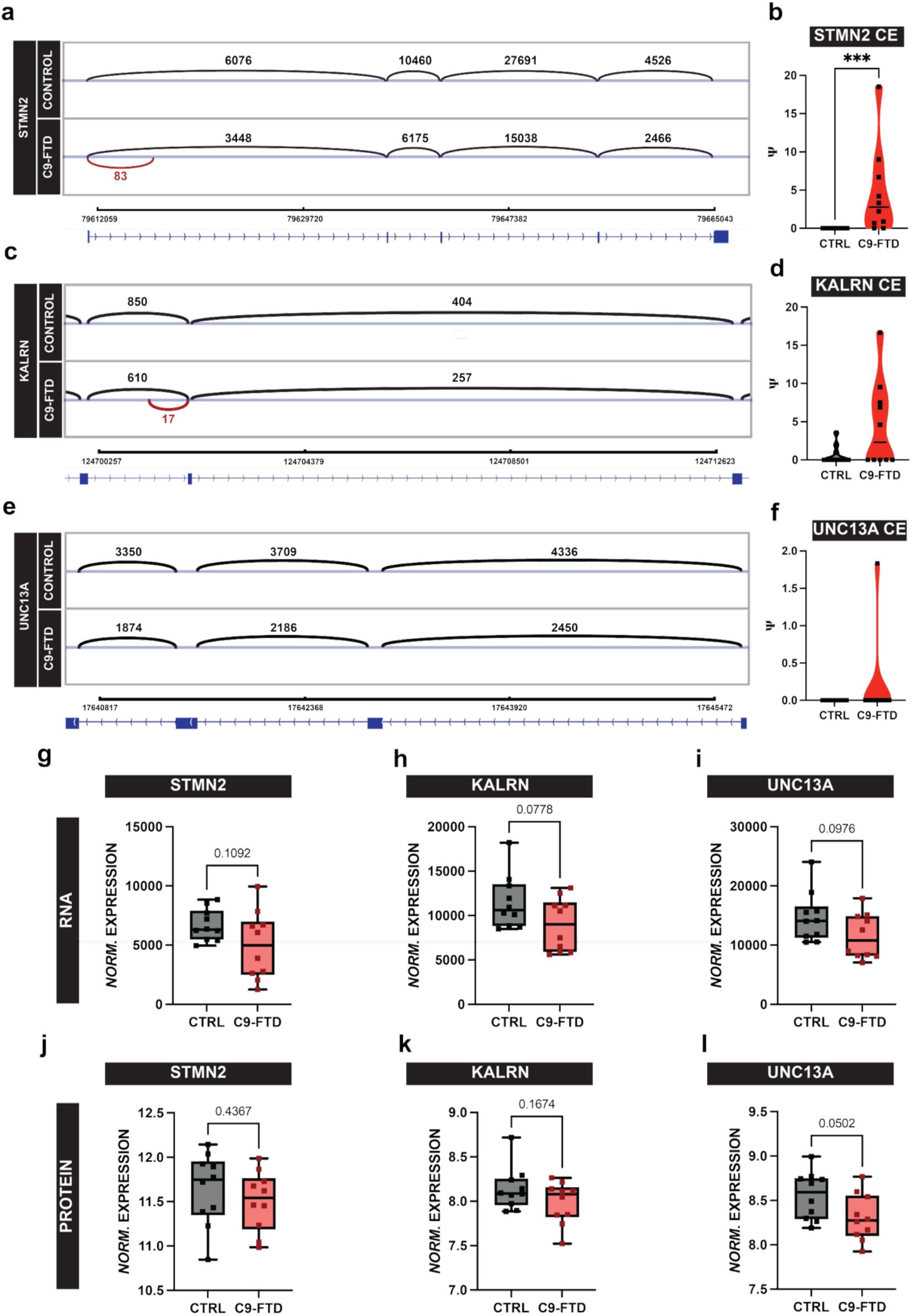
Detection of TDP-43-associated cryptic exon-containing transcripts in C9-FTD frontal cortex synaptosomes. **a: Detection of STMN2 cryptic exon-containing transcripts:** IGV sashimi plot of full-length *STMN2* gene transcripts containing cryptic exon (CE) inclusions. The top track shows the data from combined FCx control case synaptosomes (n=10). The bottom track shows the data from combined FCx C9-FTD case synaptosomes (n=10). **b: Quantification of Cryptic Exon inclusion:** Violin plot displaying the percentage spliced in (PSI) values of CE inclusion in *STMN2* within control and C9-FTD FCx synaptosomes (p=0.0007; Mann-Whitney test test). *Each dot represents each case subject used in this study.* **c:** Detection of KALRN cryptic exon-containing transcripts: IGV sashimi plot of full-length *KALRN* gene transcripts containing cryptic exon (CE) inclusions. The top track shows the data from combined FCx control case synaptosomes (n=10). The bottom track shows the data from combined FCx C9-FTD case synaptosomes (n=10). **d: Quantification of Cryptic Exon inclusion:** Violin plot displaying the PSI values of CE inclusion in *KALRN* within control and C9-FTD FCx synaptosomes (p=0.1462; Mann-Whitney test). *Each dot represents each case subject used in this study.* **e:** Detection of UNC13A cryptic exon-containing transcripts: IGV sashimi plot of full-length *UNC13A* gene transcripts containing cryptic exon (CE) inclusions. The top track shows the data from combined FCx control case synaptosomes (n=10). The bottom track shows the data from combined FCx C9-FTD case synaptosomes (n=10). **f: Quantification of Cryptic Exon inclusion:** Violin plot displaying the PSI values of CE inclusion in *UNC13A* within control and C9-FTD FCx synaptosomes (p>0.9999; Mann-Whitney test). *Each dot represents each case subject used in this study.* **g-i:** Comparison of normalized gene expression for CE-containing genes in FCx synaptosomes. *Each dot represents each case subject used in this study.* **g:** Box plot displaying normalized gene expression of *STMN2* (p=0.1092; Welch’s t-test). **h:** Box plot displaying normalized gene expression of *KALRN* (p=0.0778; Welch’s t-test). **i:** Box plot displaying normalized gene expression of *UNC13A* (p=0.0976; Welch’s t-test). **j-l:** Comparison of normalized protein expression for CE-containing genes in FCx synaptosomes. *Each dot represents each case subject used in this study*. **j:** Box plot displaying normalized protein expression of *STMN2* (p=0.4367; Welch’s t-test). **k:** Box plot displaying normalized protein expression of *KALRN* (p=0.1674; Welch’s t-test). **l:** Box plot displaying normalized protein expression of *UNC13A* (p=0.0502; Welch’s t-test).

Several limitations of the present study should be considered when interpreting these findings. First, although synaptosome preparations demonstrated robust enrichment of synaptic proteins and depletion of non-synaptic markers, the isolated fractions do not represent completely pure synaptic compartments. Residual contamination from non-synaptic cellular components is an inherent limitation of biochemical synaptosome isolation approaches and may contribute to the detection of transcripts that are not exclusively localized to synapses [22]. Consequently, while our findings demonstrate the presence of cryptic exon (CE)-containing transcripts within synaptosome-enriched fractions, they do not establish precise sub-synaptic localization. Future studies using high-resolution spatial transcriptomics, RNA in situ hybridization, or imaging-based approaches will be necessary to determine the exact localization of CE-containing transcripts within neuronal processes and synaptic compartments. A second limitation relates to the localization of CE-containing transcripts found in the synaptic compartment. Our analyses focused on previously validated TDP-43-dependent cryptic exons, including those within *STMN2*, *KALRN*, and *UNC13A* [1, 23, 31, 37, 45]. As a result, we did not perform an unbiased transcriptome-wide discovery analysis for novel cryptic exon events within synaptosomes. It is therefore possible that additional CE-containing transcripts localize to synaptic compartments and contribute to synaptic dysfunction in C9-FTD. Similarly, although we demonstrate the presence of CE-containing transcripts within synaptosome fractions, our data do not establish whether these transcripts undergo local translation or directly contribute to synaptic pathology. Determining the stability, translational competence, and functional consequences of synaptically localized aberrantly spliced transcripts represents an important area for future investigation.

Additional limitations arise from comparisons between human postmortem tissue and iPSC-cortical neuron models. Human frontal cortex tissue contains diverse neuronal and non-neuronal cell populations, including glial subtypes of astrocytes, microglia, oligodendrocytes, and vascular-associated cells, all of which influence synaptic function and disease progression [2, 8, 12, 26, 29, 48, 54, 59]. In contrast, the iPSC-derived cultures used in this study consisted of cortical neuron monocultures lacking the cellular complexity of the human brain. Given the established roles of glial cells in synaptic maturation, maintenance, and neurodegeneration-associated synaptic remodeling, differences in cellular composition likely contributed to the relatively modest overlap observed between postmortem and iPSC-derived transcriptomic and proteomic datasets. Nevertheless, both model systems demonstrated convergent alterations in pathways associated with synaptic organization, neurotransmission, and neuronal function, supporting the biological relevance of the shared findings.

Finally, the postmortem tissue analyses provide a cross-sectional view of end-stage disease and therefore cannot establish the temporal sequence of molecular events occurring during disease progression. Decreased synaptic density and progressive synapse loss are well-established pathological hallmarks of neurodegenerative disease [57] and have been documented from neuroimaging studies to postmortem studies of FTD [21–23]. The synaptosomes isolated from postmortem tissue in this study likely represent remaining and surviving synapse populations rather than synapses that have been eliminated during disease progression. Consequently, the molecular aberrations identified in this study may reflect changes of resilient or compensatory synapses. Therefore, it remains unclear whether the observed synaptic proteomic and transcriptomic alterations represent primary drivers of synaptic dysfunction, compensatory responses, or downstream consequences of neurodegeneration. Longitudinal studies using human cellular models and *in vivo* systems will be required to determine how TDP-43-dependent cryptic splicing, synaptic remodeling, and neuronal dysfunction evolve over the course of disease. Despite these limitations, the convergence of findings across independent human postmortem and patient-derived neuronal model systems provides strong evidence that synaptic RNA dysregulation, including the unexpected localization of CE-containing transcripts to synaptic compartments, represents a previously unknown feature of C9-FTD pathogenesis.

In summary, our integrated synaptosome multi-omic analyses reveal extensive molecular remodeling of C9-FTD synapses. The convergence of findings across postmortem frontal cortex tissue and patient-derived iPSC-cortical neurons supports synaptic dysfunction as an early and biologically relevant feature of C9-FTD pathogenesis. Importantly, the identification of CE-containing transcripts within synaptic compartments extends current models of TDP-43 dysfunction and a critical role of aberrantly sliced RNAs to local synaptic defects. Together, these findings establish synapses as a place of subcellular vulnerability in C9-FTD and provide a framework for future studies aimed at restoring synaptic integrity in TDP-43 associated neurodegeneration.

## SUPPLEMENT FIGURE LEGENDS

**Supplement Fig 1.**
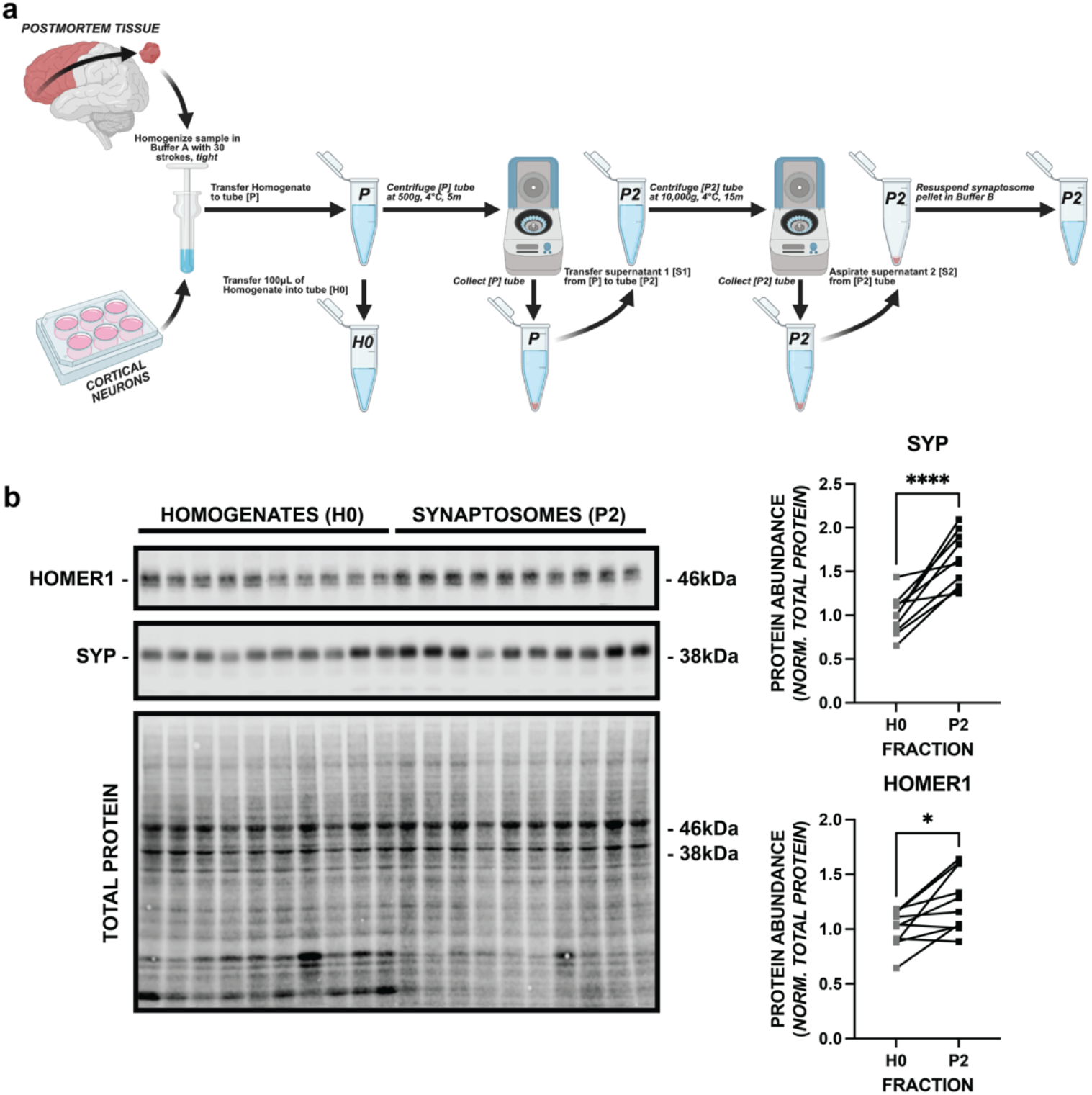
Biochemical fractionation and enrichment of frontal cortex synaptosomes. **a: Biochemical fractionation of synaptosomes:** Schematic diagram illustrating the workflow of synaptosome isolation, in both frontal cortex and iPSC-cortical neuron models, where the sample is homogenized (H0 fraction) and subjected to a series of differential centrifugation to derive the synaptosome fraction (P2) (centrifuged at 500xg to separate cytosol from cell debris, and the supernatant 1 (S1) centrifuged at 10,000xg to separate the cytosol from the crude synaptosome pellet). **b: Western Blot Validation:** Synaptic protein enrichment across homogenate (H0) and synaptosome (P2) fractions. Western blot quantification shows enrichment of presynaptic *Synaptophysin* in the P2 fraction (Welch’s t-test, p<0.0001), and enrichment of postsynaptic *Homer1* in the P2 fraction (Welch’s t-test, p<0.05), both normalized to total protein abundance.

**Supplement Fig 2.**
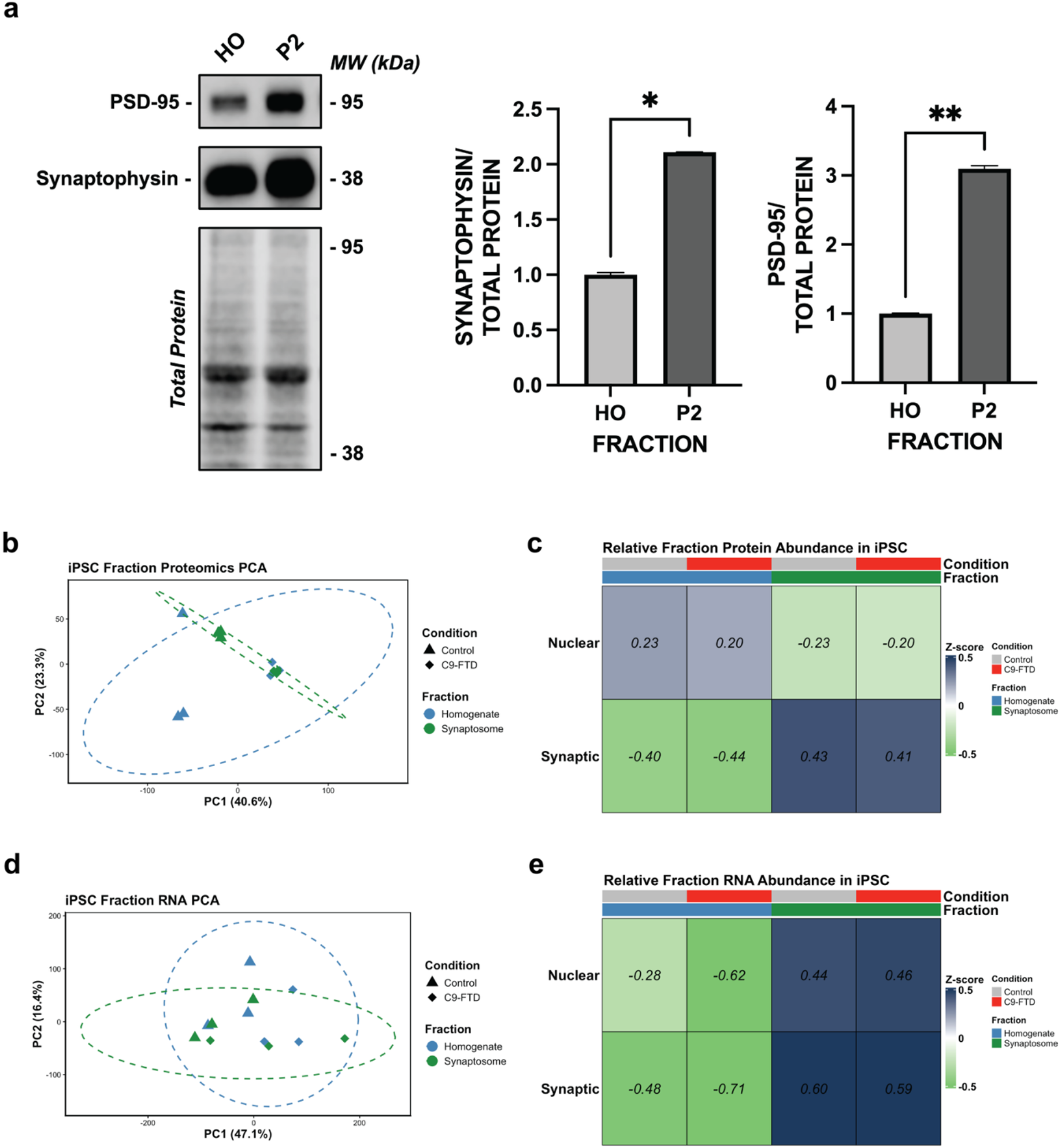
Successful synaptosome enrichment in iPSC-cortical neuron models. **a: Western Blot Validation:** Synaptic protein enrichment across homogenate (H0) and synaptosome (P2) fractions. Western blot quantification shows enrichment of presynaptic *Synaptophysin* in the P2 fraction (Welch’s t-test, p<0.05), and enrichment of postsynaptic *PSD-95* in the P2 fraction (Welch’s t-test, p<0.05), both normalized to total protein abundance. **b: Distinct Molecular Profiles by Fraction:** Principal Component Analysis (PCA) of proteomic data from homogenate (H0) and synaptosome (P2) fractions across control and C9-FTD iPSC-cortical neurons (PCA1=40.6%, PC2=23.3%), demonstrating sample separation by disease status and biochemical fraction identity. **c: Compartmental Enrichment of Synaptic Signatures:** Z-score-normalized heatmap of synaptic protein abundance across H0 and P2 fractions in control and C9-FTD iPSC-cortical neurons, highlighting enrichment of synaptic gene sets in the P2 fraction and nuclear gene set enrichment in the H0 fraction. **d: Distinct Molecular Profiles by Fraction:** PCA of transcriptomic data from iPSC-cortical neuron-derived H0 and P2 fractions (PC1=47.1%, PC2=16.4%), demonstrating sample separation by disease status and biochemical fraction identity. **e: Compartmental Enrichment of Synaptic Signatures:** Z-score-normalized heatmap of synaptic RNA abundance across H0 and P2 fractions, highlighting enrichment of synaptic gene sets in the P2 fraction. **c, e) Fraction Purity Validation:** Heatmaps showing Z-score normalized abundance of synaptic and nuclear markers. The Synaptosome fraction shows a strong positive enrichment of synaptic markers compared to the Homogenate. Conversely, nuclear markers are depleted in the Synaptosome fraction relative to the Homogenate, confirming successful separation of cytoplasmic synaptic terminals from nuclear contaminants.

**Supplement Fig 3.**
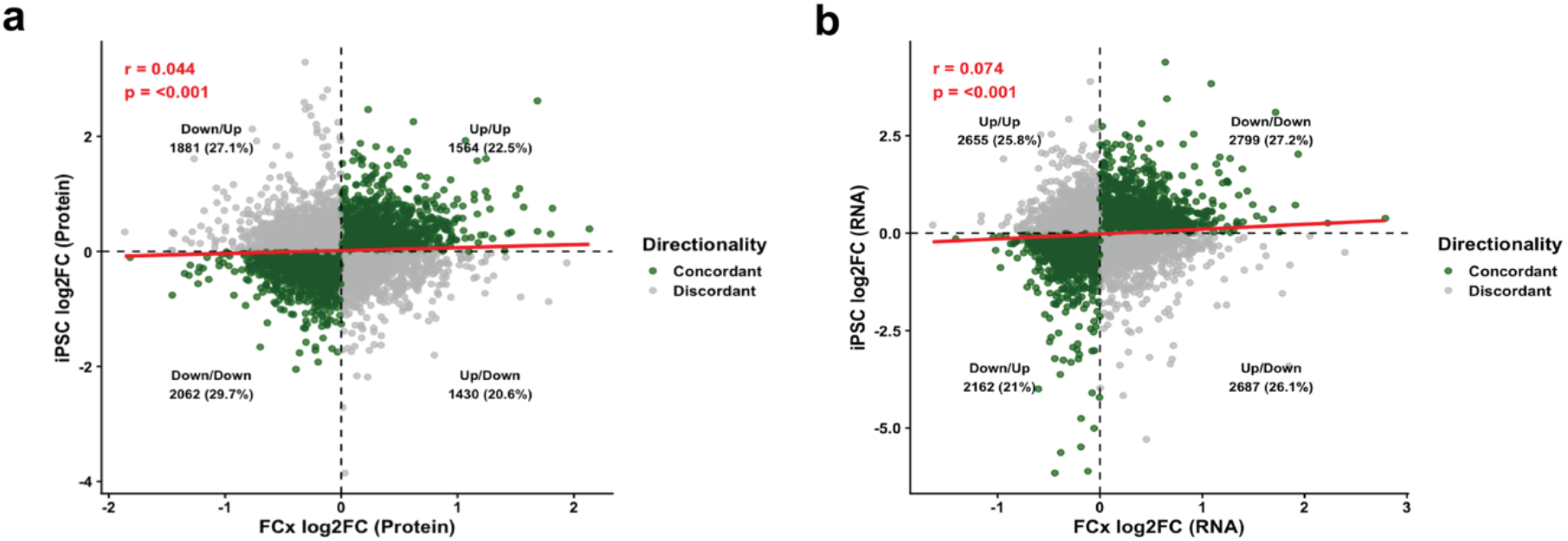
Correlation of protein and RNA expression across frontal cortex and iPSC-derived synaptosomes. **a: Protein Concordance across Frontal Cortex and iPSCs:** Correlation analysis of frontal cortex synaptic proteome (x-axis) and iPSC synaptosome proteome (y-axis) log_2_FC. The correlation was modest (Pearson r=0.044, p<0.001), however a substantial proportion of proteins exhibited concordant regulation across model systems, with 22.5% of proteins concordantly upregulated and 29.7% of proteins concordantly downregulated. The presence of discordant proteins (grey) highlights specific differences between frontal cortex and iPSC-derived synaptic proteomes. **b: Transcriptional Concordance across Frontal Cortex and iPSCs:** Correlation analysis of frontal cortex synaptosomal transcriptome (x-axis) and iPSC synapse transcriptome (y-axis) log_2_FC. The correlation was modest (Pearson r=0.074, p<0.001) and exceeded proteomic observations. 27.2% of transcripts were concordantly upregulated and 21% of transcripts were concordantly downregulated across model systems, with the presence of discordant transcripts (grey) between models. These changes reflect shared and model-specific aberrations.

**Figure 4.**
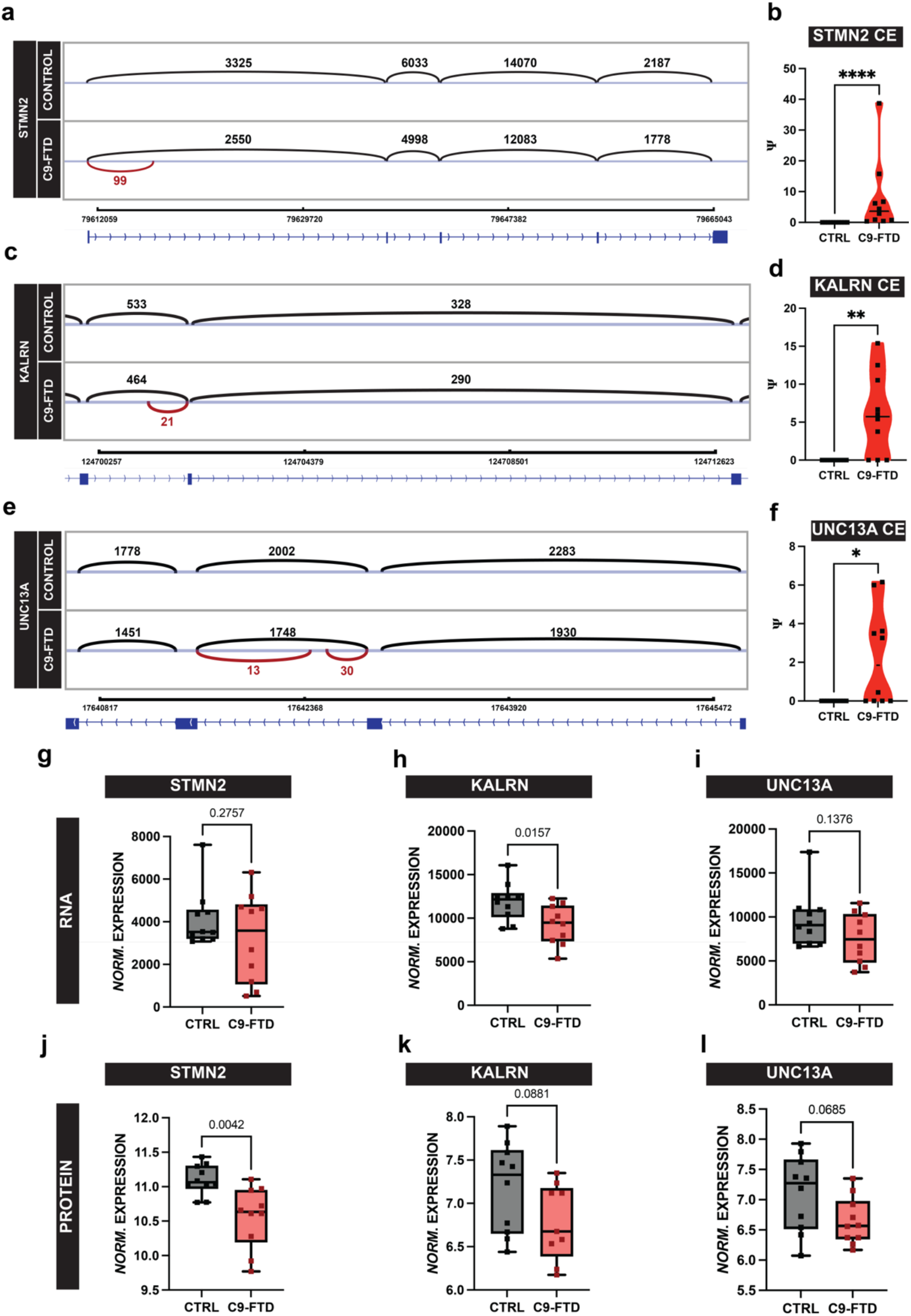
Alternative Splicing analysis reveals cryptic exon inclusion in C9-FTD frontal cortex homogenates. **a: Detection of STMN2 cryptic exon-containing transcripts:** IGV sashimi plot of full-length *STMN2* gene transcripts containing cryptic exon (CE) inclusions. The top track shows the data from combined FCx control cases (n=10). The bottom track shows the data from combined FCx C9-FTD cases (n=10). **b: Detection of Cryptic Exon inclusion:** Violin plot displaying the percentage spliced in (PSI) values of CE inclusion in *STMN2* within control and C9-FTD FCx homogenates (p<0.0001; Mann-Whitney test test). *Each dot represents each case subject used in this study.* **c:** Detection of KALRN cryptic exon-containing transcripts: IGV sashimi plot of full-length *KALRN* gene transcripts containing cryptic exon (CE) inclusions. The top track shows the data from combined FCx control cases (n=10). The bottom track shows the data from combined FCx C9-FTD cases (n=10). **d: Quantification of Cryptic Exon inclusion:** Violin plot displaying the PSI values of CE inclusion in *KALRN* within control and C9-FTD FCx homogenates (p=0.0031; Mann-Whitney test). *Each dot represents each case subject used in this study.* **e:** Detection of UNC13A cryptic exon-containing transcripts: IGV sashimi plot of full-length *UNC13A* gene transcripts containing cryptic exon (CE) inclusions. The top track shows the data from combined FCx control cases (n=10). The bottom track shows the data from combined FCx C9-FTD cases (n=10). **f: Quantification of Cryptic Exon inclusion:** Violin plot displaying the PSI values of CE inclusion in *UNC13A* within control and C9-FTD FCx homogenates (p=0.0108; Mann-Whitney test). *Each dot represents each case subject used in this study.* **g-i:** Comparison of normalized gene expression for CE-containing genes in FCx homogenates. *Each dot represents each case subject used in this study.* **g:** Box plot displaying normalized gene expression of *STMN2* (p=0.2757; Welch’s t-test). **h:** Box plot displaying normalized gene expression of *KALRN* (p=0.0157; Welch’s t-test). **i:** Box plot displaying normalized gene expression of *UNC13A* (p=0.1376; Welch’s t-test). **j-l:** Comparison of normalized protein expression for CE-containing genes in FCx homogenates. *Each dot represents each case subject used in this study.* **j:** Box plot displaying normalized protein expression of *STMN2* (p=0.0042; Welch’s t-test). **k:** Box plot displaying normalized protein expression of *KALRN* (p=0.0881; Welch’s t-test). **l:** Box plot displaying normalized protein expression of *UNC13A* (p=0.0685; Welch’s t-test).

## Supporting information

Supplement Table 1: Postmortem Tissue Demographics

Supplement Table 2: Antibodies used in this study

## MANUSCRIPT ACKNOWLEDGEMENTS

The authors wish to thank University College London Queen Square Brain Bank for Neurological Disorders (London, UK) for the provision of human postmortem tissue used in this study. The Queen Square Brain Bank is supported by the Reta Lila Weston Institute of Neurological Studies, UCL Queen Square Institute of Neurology. Additionally, the authors thank Prof. Selina Wray, PhD, for the provision of the patient-derived cell lines used for cortical neuron differentiation used in this study, from University College London Queen Square Institute of Neurology. We thank Zhuo Li, PhD, in the Electron Microscopy Core at City of Hope (Duarte, California). We thank the Integrated Mass Spectrometry Shared Resource at TGEN for the processing of our protein samples. We thank Dominic Julian, PhD, at Barrow Neurological Institute for critical discussions of the datasets.

## AUTHOR CONTRIBUTIONS

AMS and RGS conceptualized the study; AMS, RGS, LMG, KVK-J, PP provided oversight over the execution of the study; AMS, LMG, KP performed wet lab experiments; RS and MM performed the sample preparation for mass spectrometry; AB performed the sample preparation for RNA sequencing; EBA, KG-M, IP, and MH performed bioinformatics analysis; AMS, EBA, and KG-M performed data analysis and interpretation. AMS and RGS wrote the manuscript with input from all co-authors.

*AMS: Ashton M Spillman; EBA: Eric B. Alsop; LMG: Lauren M. Gittings; KG-M: Krystine Garcia-Mansfield; IP: Ignazio Piras; AB: Anna Bonfitto; MM: Melissa Martinez; RS: Ritin Sharma; KP: Kim R. Preller; MH: Matthew Huentelman; PP: Patrick Pirrotte; KVK-J: Kendall Van Keuren-Jensen; RGS: Rita Sattler*

