## Supplementary figures and images for "Integrative synaptosome multi-omics reveals disrupted synapse organization and localized cryptic transcripts in *C9ORF72*-Frontotemporal Dementia"

### Supplement Table 1: Postmortem Tissue Demographics

**SUPPLEMENT TABLES**

**Supplement Table 1. Frontal Cortex Tissue Demographics.**

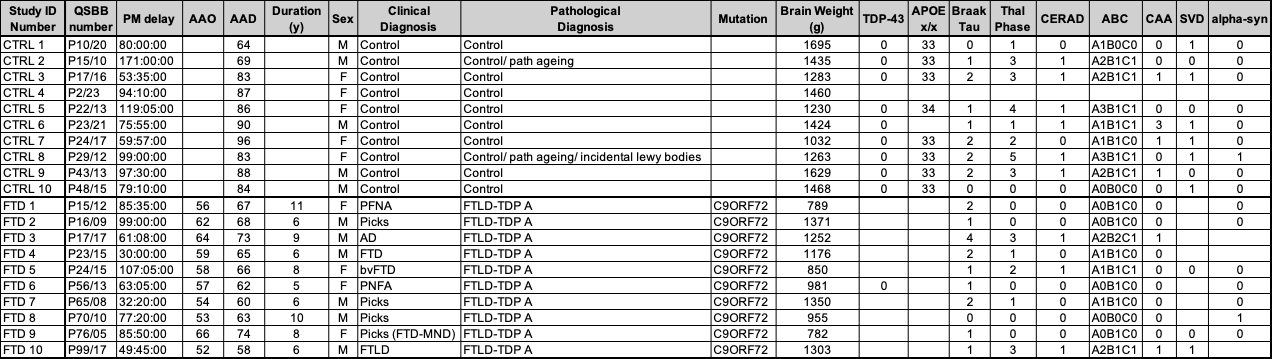
