## Supplement Table 2: Antibodies used in this study for "Integrative synaptosome multi-omics reveals disrupted synapse organization and localized cryptic transcripts in *C9ORF72*-Frontotemporal Dementia"

| **Application** | **Primary Antibody** | **Species** | **Dilution** | **1° Catalog #** | **Secondary Antibody** | **Dilution** | **2° Catalog #** |
| --- | --- | --- | --- | --- | --- | --- | --- |
| Immunocytochemistry | VGLUT1 | Rb | 1:1000 | Synaptic Systems #135303 | Goat anti-Rabbit Secondary Antibody, Alexa Fluor™ 488 | 1:1000 | Invitrogen, #A-11034 |
|  | VGAT | Rb | 1:1000 | Synaptic Systems #131002 |  |  |  |
|  | Homer1 | Ch | 1:500 | Synaptic Systems #160006 | Goat anti-Chicken Secondary Antibody, Alexa Fluor™ 555 | 1:1000 | Invitrogen, #A-21437 |
|  | Gephyrin | Ch | 1:1000 | Synaptic Systems #147009 |  |  |  |
|  | MAP2 | GP | 1:1000 | Synaptic Systems #188004 | Goat anti-Guinea Pig Secondary Antibody, Alexa Fluor™ 633 | 1:1000 | Invitrogen, #A-21105 |
| Western Blot | Synaptophysin | Rb | 1:200 | Abcam #16659 | HRP, Donkey anti-Rabbit IgG (H+L) Secondary Antibody | 1:10,000 | Invitrogen, #31458 |
|  | Homer1 | Rb | 1:500 | Abcam #184955 |  |  |  |
|  | PSD-95 | Rb | 1:500 | Abcam #18258 |  |  |  |
